# A Cryptochrome adopts distinct moon- and sunlight states and functions as sun- versus moonlight interpreter in monthly oscillator entrainment

**DOI:** 10.1101/2021.04.16.439809

**Authors:** Birgit Poehn, Shruthi Krishnan, Martin Zurl, Aida Coric, Dunja Rokvic, N. Sören Häfker, Enrique Arboleda, Lukas Orel, Florian Raible, Eva Wolf, Kristin Tessmar-Raible

**Author notes:** correspondence: Eva Wolf, Kristin Tessmar-Raible. equal contribution (Birgit Poehn, Shruthi Krishnan, Aida Coric, Dunja Rokvic).

## Abstract

The moon’s monthly cycle synchronizes reproduction in countless marine organisms. The mass-spawning bristle worm *Platynereis dumerilii* uses an endogenous monthly oscillator to phase reproduction to specific days. Classical work showed that this oscillator is set by full moon. But how do organisms recognize such a specific moon phase? We uncover that the light receptor L-Cryptochrome (L-Cry) is able to discriminate between different moonlight durations, as well as between sun- and moonlight. Consistent with L-Cry’s function as light valence interpreter, its genetic loss leads to a faster re-entrainment under artificially strong nocturnal light. This suggests that L-Cry blocks “wrong” light from impacting on the monthly oscillator. A biochemical characterization of purified L-Cry protein, exposed to naturalistic sun- or moonlight, reveals the formation of distinct sun- and moonlight states characterized by different photoreduction- and recovery kinetics of L-Cry’s co-factor Flavin Adenine Dinucleotide. *In vivo,* L-Cry’s sun-versus moonlight states correlate with distinct sub-cellular localizations, indicating different signalling. In contrast, r-Opsin1, the most abundant ocular opsin, is not required for monthly oscillator entrainment. Our work reveals a new concept for correct moonlight interpretation involving a “valence interpreter” that provides entraining photoreceptor(s) with light source and moon phase information. These findings advance our mechanistic understanding of a fundamental biological phenomenon: moon-controlled monthly timing. Such level of understanding is also an essential prerequisite to tackle anthropogenic threats on marine ecology.

## Main text

Lunar influences on animals are especially well documented in the marine environment ^1–3^. Starting with the early 20th century, numerous scientific studies have shown that the reproductive behavior and sexual maturation of animals as diverse as corals, polychaetes, echinoderms, fishes or turtles are synchronized by the lunar cycle ^1, 3–7^. In addition, lunar timing effects have also been documented outside the marine environment ^8, 9^, and recently uncovered correlations of human sleep and menstrual cycle properties with moon phases have re-initiated the discussion of an impact of the moon even on human biology ^10, 11^. As recently documented for corals, desynchronization of these reproductively critical rhythms by anthropogenic impacts poses a threat to species survival ^12^.

Despite the importance and widespread occurrence of lunar rhythms, functional mechanistic insight is lacking. Importantly, this synchronization among conspecifics is in many cases not simply a direct reaction to a stimulus, but instead governed by endogenous monthly oscillators: circalunar clocks ^3, 9, 13–16^. The marine bristle worm *Platynereis dumerilii* is well-documented to possess such a circalunar clock, which controls its reproductive timing and can be entrained by nocturnal light in the lab ^5, 15, 17^. Several reports have linked the expression of cryptochromes (CRYs) with moon phase, suggesting that these genes could be involved in circalunar time-keeping ^18^, possibly- as proposed for corals- as lunar light receptors ^2, 19, 20^. However, no functional molecular support for such an involvement exists. In order to move from expression correlation to a mechanistic understanding, we investigated the functional role and biochemical properties of the light-receptive cryptochrome L-Cry in the annelid *Platynereis dumerilii* ^15^.

In order for organisms to synchronize their monthly oscillator they need to be entrained to a specific moon phase. However, this requires that the entraining mechanisms can discriminate between naturalistic sun- and moonlight, as well as the different moon phases. The latter differ by moonlight intensity and the duration^21^. Classical and recent work on the monthly oscillators of the midge *Clunio* and the bristle worm *Platynereis* established that mimicking the duration of full moon (i.e. continuous light at night) is sufficient to entrain their monthly oscillators ^5, 13, 15, 22^. We thus followed and subsequently expanded on these established experimental paradigms and focussed on new moon versus full moon conditions throughout the experiments.

Prompted by the clear prediction from corals ^2^ and our own previous study showing that *Pdu-*L-Cry functions in principle as a light receptor in a heterologous cell expression system ^15^, we studied genetic loss-of-function mutants of *Platynereis l-cry* and combined this work with biochemical and *in vivo* cell biological analyses using custom-designed light sources based on long-term measurements of light intensity and spectra at the worms’ natural habitat.

### *l-cry* mutants show higher spawning synchrony than wild-type animals under non-natural light conditions

In order to test for a functional involvement of L-Cry in monthly oscillator function, we generated two *l-cry* mutant alleles (Δ34 and Δ11bp) (Fig.1a) using TALENs ^23^. In parallel, we generated a monoclonal antibody against *Platynereis* L-Cry. By testing mutant versus wildtype worms with the anti-L-Cry antibody in Western blots (Fig.1b) and immunohistochemistry (Fig. 1e-j), we verified the absence of L-Cry protein in mutants. Furthermore, we confirmed that the staining of the antibody in wildtype worms (Fig. 1e-h) matches the regions where *l-cry* mRNA is expressed (Fig. 1d). These tests confirmed that the engineered *l-cry* mutations result in loss-of-function alleles. In turn, they validate the specificity of the raised anti-L-Cry antibody.

**Figure 1.**
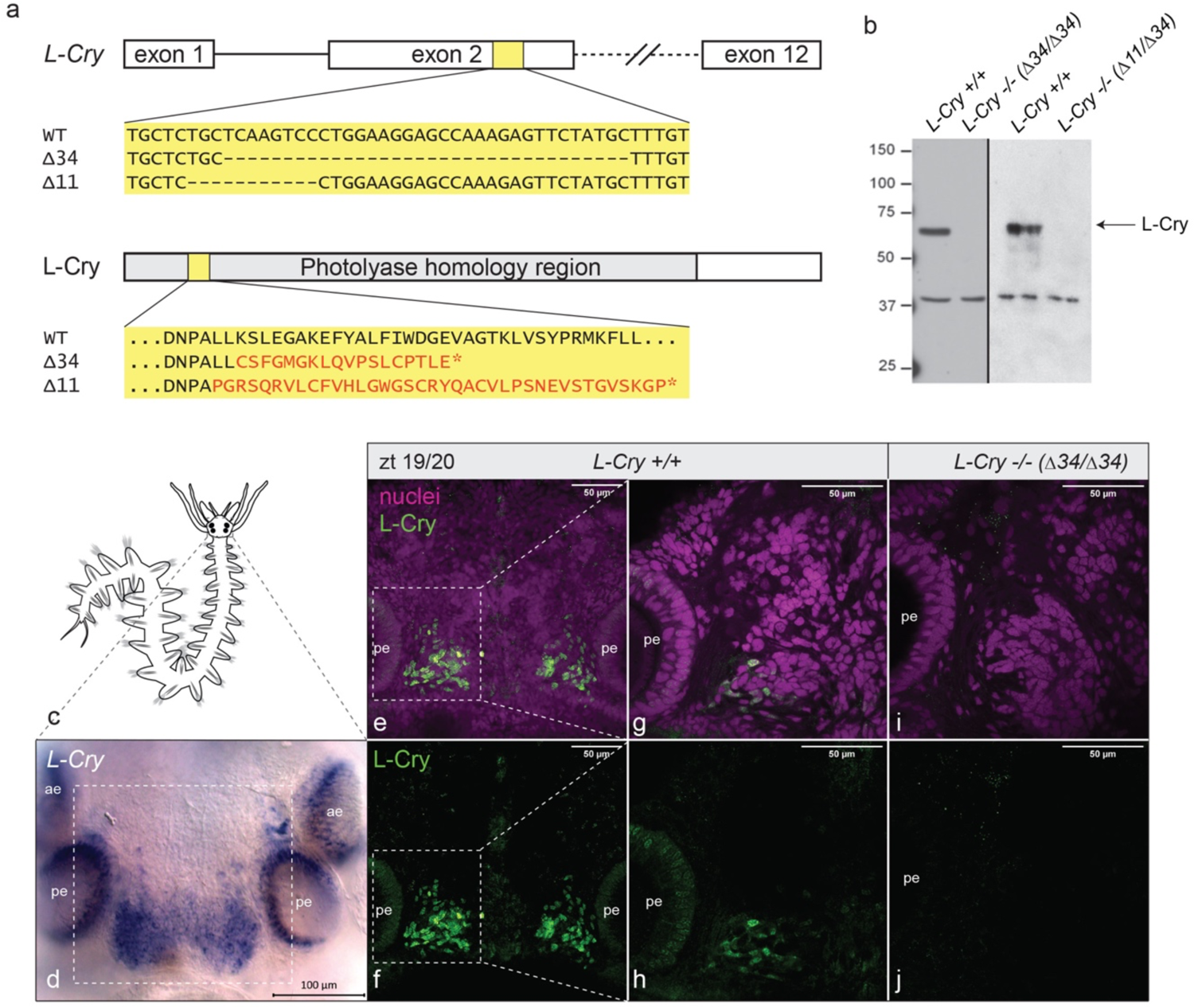
*l-cry ^-/-^* mutants are loss-of-function alleles. **(a)** Scheme of the *l-cry* genomic locus for wt and mutants. Both mutant alleles result in an early frameshift and premature stop codons. The Δ34 allele has an additional 9bp deletion in exon 3. **(b)** Western Blots of *P. dumerilii* heads probed with anti-L-Cry antibody. **(c)** scheme of *P. dumerilii.* **(d)** whole mount *in situ* hybridization against *l-cry* mRNA on worm head. ae, anterior eye; pe, posterior eye. **(e-j)** Immunohistochemistry of premature wildtype (e-h) and mutant (i,j) worm heads sampled at zt19/20 using anti-L-Cry antibody (green) and Hoechst staining (magenta), dorsal views, anterior up. e,f: z-stack images (maximal projections of 50 layers, 1.28µm each) in the area highlighted by the rectangle in (d), whereas (g-j) are single layer images of the area highlighted by the white rectangles in (e) and (f).

We next assessed the circalunar maturation timing of wildtypes and *l-cry* mutant populations in conventional culture conditions, i.e. worms grown under typical indoor room lighting (named here artificial sun- and moonlight, Extended Data Figure 1b).

We expected either no phenotype (if L-Cry was not involved in circalunar clock entrainment) or a decreased spawning precision (if L-Cry was functioning as moonlight receptor in circalunar clock entrainment). Instead we observed an increased precision of the entrained worm population: We analysed the maturation data using two statistical approaches, linear and circular statistics. We used the classical linear plots ^5^ and statistics to compare the monthly spawning data distribution (Fig2a-c,i). This revealed a clear difference between mutant animals, which exhibited a stronger spawning peak at the beginning of the NM phase, compared to their wildtype and heterozygous counterparts (Fig. 2a-c, Kolmogorov-Smirnov-Test on overall data distribution, Fig. 2i).

**Figure 2:**
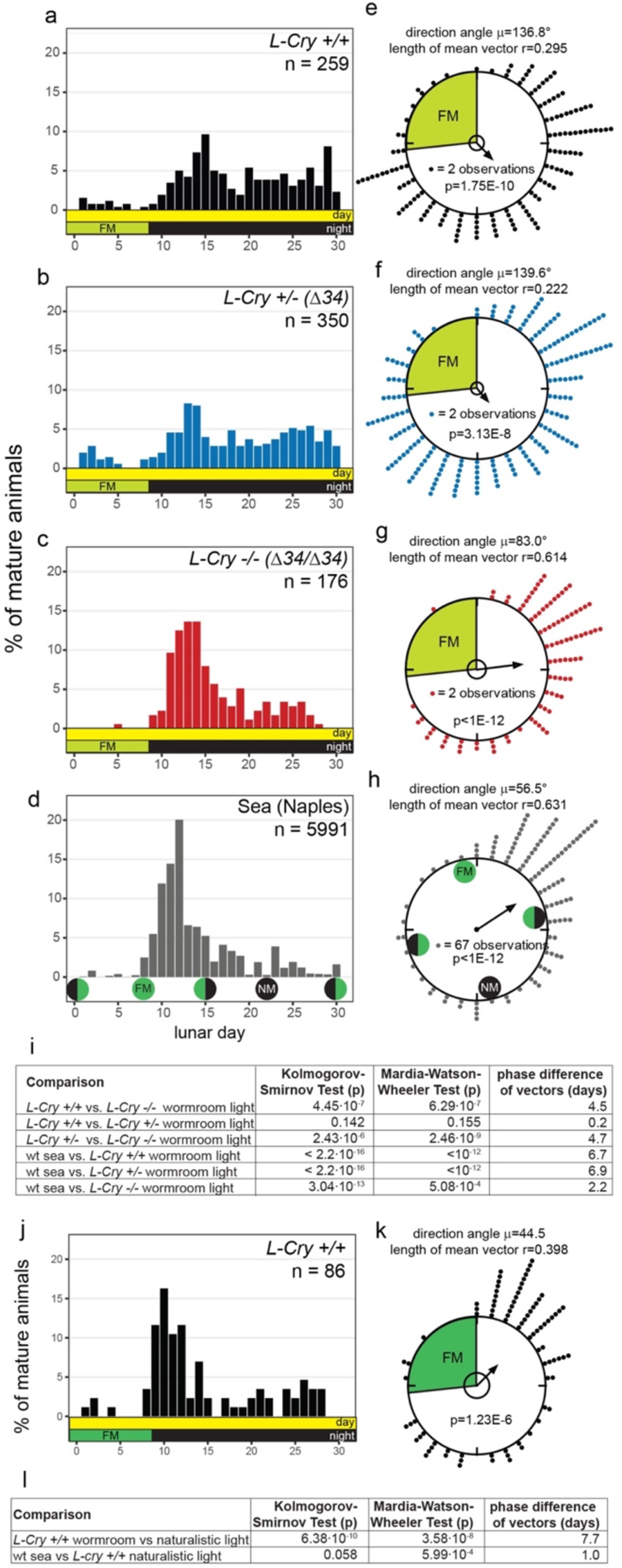
L-Cry shields the circalunar clock from light that is not naturalistic moonlight. (**a-d,j)** Spawning of *l-cry* ^+/+^ (a), *l-cry* ^+/– (Δ34)^ **(a) (b)** and *l-cry* ^-/-(Δ34/ Δ34)^ (c) animals over the lunar month in the lab with 8 nights of artificial moonlight (a-c), under natural conditions in the sea (d, replotted from ref. ^29,31^) and in the lab using naturalistic sun- and moonlight (j, 8 nights moonlight). **(e-h,k)** Data as in (a-d,j) as circular plot. 360° correspond to 30 days of the lunar month. The arrow represents the mean vector, characterized by the direction angle µ and r (length of µ). r indicates phase coherence (measure of population synchrony). p-values inside the plots: result of Rayleigh Tests. Significance indicates non-random distribution of data points. The inner circle represents the Rayleigh critical value (p=0.05). **(i,l)** Results of multisample statistics on spawning data shown in (a-h,j,k). The phase differences in days can be calculated from the angle between the two mean vectors (i.e. 12°= 1 day).

We then analysed the same data using circular statistics (as the monthly cycle is repeating, see details in Methods section), which allowed us to describe the data with the mean vector (defined by the direction angle µ and its length r, shown as arrows in Fig. 2e-g). The phase coherence r (ranging from 0 to 1) serves as a measure for synchrony of the population data. As expected for entrained populations, all genotypes distributed their spawning across a lunar month significantly different from random (Fig. 2e-g, p-values in circles, Rayleigh’s Uniformity test ^24^). In line with the observed higher spawning peak of the *l-cry-/-* mutants in the linear plots, the circular analysis revealed a significant difference in spawning distribution (Mardia-Watson-Wheeler test, for details see Methods section) and higher spawning synchrony of mutants (r=0.614) than in wildtypes and heterozygotes (r=0.295 and r=0.222) (Fig. 2i). The specificity of this phenotype of higher spawning precision for *l-cry* homozygous mutants was confirmed by analyses on trans-heterozygous *l-cry* (Δ34/Δ11) mutants (Extended Data Figure 2), and by the fact that such a phenotype is not detectable in any other light receptor mutant available in *Platynereis* (*r-opsin1* ^25^: Extended Data Figure 3a,b,e,f,i; *c-opsin1* ^26^: Extended Data Figure 3c,d,g,h,i, *Go-opsin*: refs. ^27, 28^).

### The higher spawning synchrony of *l-cry* mutants under artificial light mimics the spawning precision of wild-type at its natural habitat

This increased spawning precision of *l-cry* mutants under artificial (but conventional indoor) laboratory light conditions let us wonder about the actual population synchrony of the worms under truly natural conditions. The lunar spawning synchrony of *Platynereis dumerilii* at the Bay of Naples (the origin of our lab culture) has been worked on for more than 100 years. This allowed us to re-investigate very detailed spawning data records from the worms’ natural habitat published prior to environmental/light pollution. For better accessibility and comparability we combined all months and replotted the data published in 1929 ^29^ (Fig. 2d,h,I; see details in Methods section; r=0.631). This analysis revealed that the higher spawning synchrony in *l-cry*^-/-^ worms mimics the actual spawning synchrony of *Platynereis dumerillii* populations in their natural habitat ^29^ (compare Fig.2c,g with 2d,h.)

Given that recent, non-inbred isolates from the same habitat as our lab inbred strains (which is the same habitat as the data collected in ref. ^29^) exhibit a broad spawning distribution under standard worm culture light conditions (which includes the bright artificial moon light) ^30^, we hypothesized that the difference in spawning synchrony between wildtype laboratory cultures and populations in their natural habitat is caused by the rather bright nocturnal light stimulus typically used for the standard laboratory culture (Extended Data Figure 1a vs. b).

### Lunar spawning precision of wild-type animals depends on naturalistic moonlight conditions

We next tested the resulting prediction that naturalistic moonlight should increase the spawning precision of the wildtype population, using naturalistic sun- and moonlight devices we specifically designed based on light measurements at the natural habitat of *Platynereis dumerilii* ^26^ (Extended Data Extended Data Fig. 1a,c). We assessed the impact of the naturalistic sun- and moonlight (Extended Data Fig. 1a,c) on wildtype animals, maintaining the temporal aspects of the lab light regime (i.e. 8 nights of “full moon”). Indeed, merely adjusting the light intensity to naturalistic conditions increased the precision and phase coherence of population-wide reproduction: After several months under naturalistic sun- and moonlight, wildtype worms spawned with a major peak highly comparable to the wildtype precision reported at its natural habitat (Fig. 2d,h vs. j,k), and also exhibited an increased population synchrony (r=0.398 compared to r=0.295 under standard worm room light conditions). This increased similarity to the spawning distribution at the natural habitat (“Sea”) is confirmed by statistical analyses (Fig. 2l): The phase difference (angle between the two mean vectors) is only one day (corresponding to 12°). In contrast, the spawning distribution of wildtypes under standard worm room light versus naturalistic light conditions is highly significantly different in linear and circular statistical tests and has a phase difference of 7.7 days (Fig. 2l).

These findings show that it is the naturalistic light that is critical for a highly precise entrainment of the monthly clock of wildtype worms. Given that *l-cry^-/-^* animals reach this high precision with the artificial light (i.e. standard lab light) implies that in wildtype L-Cry blocks artificial, but not naturalistic full moon light from efficiently synchronizing the circalunar clock. This block is removed in *l-cry^-/-^* animals, leading to a better synchronization of the *l-cry^-/-^* population. This finding suggests that L-Cry’s major role could be that of a gate-keeper controlling which ambient light is interpreted as full moon light stimulus for circalunar clock entrainment.

### *l-cry* functions as a light signal gate-keeper for circalunar clock entrainment

A prediction of this hypothesis is that mutants should entrain better to an artificial full moon light stimulus provided out-of-phase than their wildtypes counterparts (in which L-Cry should block the “wrong” moonlight at least partially from re-entraining the circalunar oscillator).

We thus compared the spawning rhythms of *l-cry*^+/+^ and *l-cry*^-/-^ worms under a re-entrainment paradigm, where we provided our bright artificial culture full moon light at the time of the subjective new moon phase (scheme Fig.3a). In order to compare the spawning data distribution relative to the initial full moon (FM) stimulus, as well as to the new full moon stimulus (i.e. new FM), we used two nomenclatures for the months: months with numbers are analyzed relative to the initial nocturnal light stimulus (i.e. FM), whereas months with letters are analyzed relative to the new (phase-shifted) nocturnal light stimulus (i.e. new FM, Fig.3a). When the nocturnal light stimulus is omitted (to test for the oscillator function) we then refer to ‘free-running FM’ (FR-FM) or ‘new free-running FM’ (new FR-FM), respectively (Fig.3a). Using these definitions, the efficiency of circalunar clock re-entrainment will be reflected in the similarity of spawning data distributions between month 1 and month D, i.e. the more similar the distribution, the more the population has shifted to the new phase.

When using the artificial nocturnal light conditions, the re-entrainment of *l-cry*^-/-^ animals was both faster and more complete than for their wildtype relatives, as predicted from our gate keeper hypothesis. This is evident from the linear data analysis and Kolmogorov-Smirnov tests when comparing the month before the entrainment (month 1) with two months that should be shifted after the entrainment (months C,D, Fig. 3b,c,f,g).

**Figure 3:**
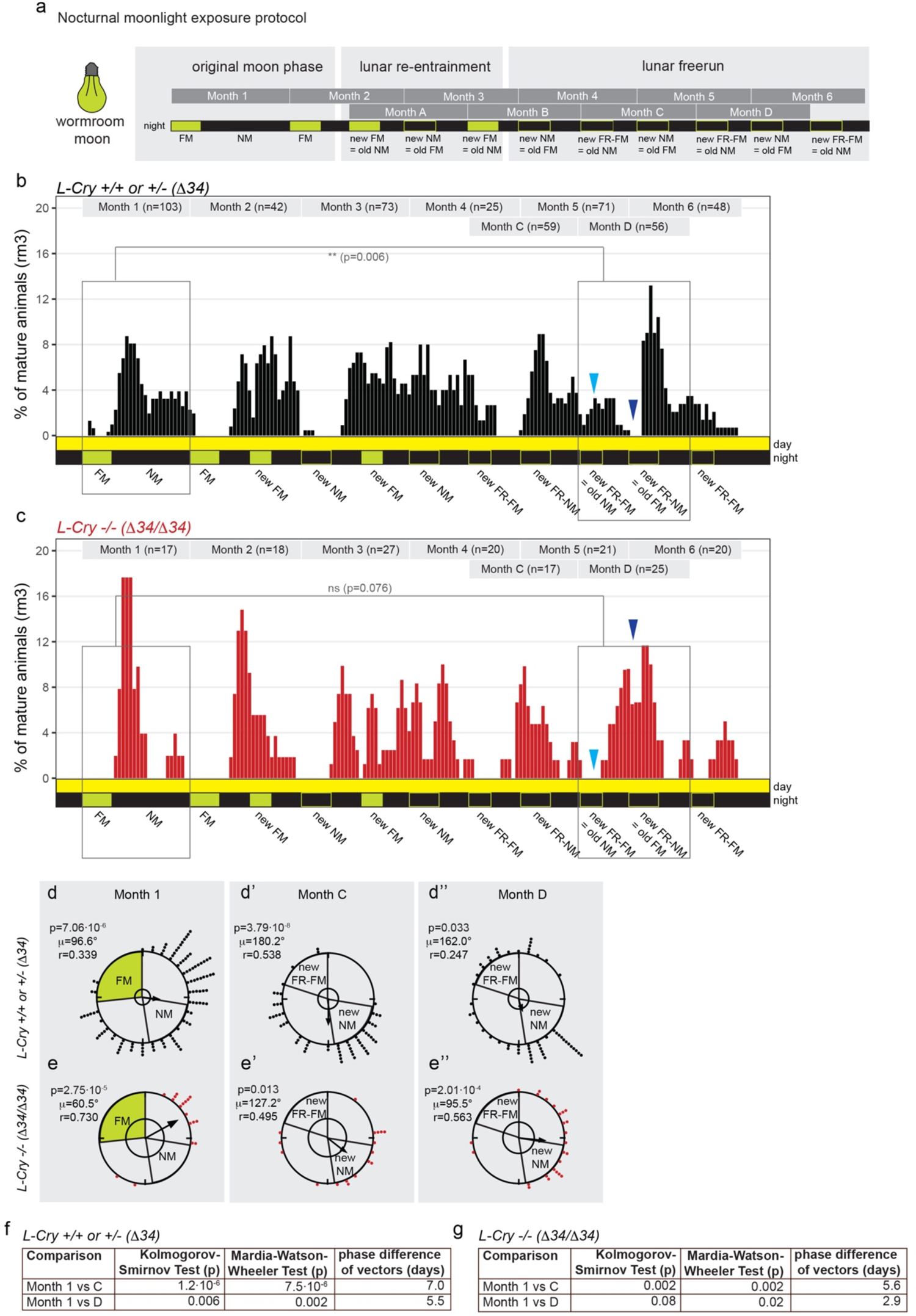
*l-cry-/-* mutants entrain the circalunar clock faster than wt to a high intensity artificial moonlight stimulus. **(a)** Nocturnal moonlight exposure protocol of lunar phase shift (entrained by 8 nights, phased shifted by 6 nights of artificial culture moon, light green). **(b,c)** Number of mature animals (percent per month, rolling mean with a window of 3 days) of *l-cry* wildtype (b) and homozygous mutant (c) animals. p-values indicate results of Kolomogorov-Smirnov tests. Dark blue arrowheads-old FM phase: wt show a spawning minimum, indicative that the worms are not properly phase shifted. Mutants spawn in high numbers, but don’t spawn at the old NM indicated by light blue arrowhead. Also compare to initial FM and NM in months 1,2. **(d-e)** Circular plots of the data shown in (b) and (c). Each circle represents one lunar month. Each dot represents one mature worm. The arrow represents the mean vector characterized by the direction angle µ and r. r (length of µ) indicates phase coherence (measure of population synchrony). The inner circle represents the Rayleigh critical value (p=0.05). **(f,g)** Results of multisample statistics of data in (d,e). Phase differences in days can be calculated from the angle between the two mean vectors (i.e. 12°= 1 day).

Most notably, while *l-cry-/-* worms were fully shifted in month D (Fig.3c: compare boxes and see complete lack of spawning at the light blue arrowhead indicating the old NM/new FR-FM phase versus massive spawning at new NM phase around dark blue arrowhead), wildtype animals were still mostly spawning according to the initial lunar phase (Fig.3b: compare boxes and see spawning at the light blue arrowhead versus almost lack of spawning at dark blue arrowhead). The faster re-entrainment of *l-cry*^-/-^, compared to *l-cry+/+* animals is also confirmed by the Mardia-Watson-Wheeler test (see Methods section for details), which shows less/no significance in the comparison of mutants before and after entrainment, but very highly significant differences in the distribution of the wildtype spawning data in the same comparison. Consistently, the phase differences in days calculated from the angle between the two mean vectors from the circular analysis is smaller in the mutants than in the wildtypes when comparing the phase of the month before the entrainment (month 1) with two months after the entrainment (months C,D) (Fig. 3d-g). The fact that there are still differences in the mutant population before and after entrainment is likely due to the fact that even the mutants are not fully re-entrained. However, they have significantly shifted stronger in response to an artificial nocturnal light stimulus than the wildtypes. This provides further evidence that in wildtype worms L-Cry indeed blocks the “wrong” light from entering into the circalunar clock and thus functions as a light gate-keeper.

### L-Cry functions mainly as light interpreter, while its contribution as direct moonlight entraining photoreceptor is minor

We next tested to which extent L-Cry function itself as a sensor for re-entrainment signal under naturalistic light conditions. Based on the finding that *l-cry-/-* worms can still re-entrain the circalunar oscillator (see above), it is clear that even if L-Cry also directly contributed to the entrainment, it cannot be the only moonlight receptor mediating entrainment. With the experiments below, we aimed to test if L-Cry has any role as an entraining photoreceptor to the monthly oscillator.

Thus, we tested how the circalunar clock is shifted in response to a re-entrainment with naturalistic moonlight in *Platynereis* wt versus *l-cry-/-* worms. For this, animals initially raised and entrained under standard worm room light conditions of artificial sun- and moonlight (Extended Data Figure 1b,e) were challenged by a deviating FM stimulus of 8 nights of naturalistic moonlight (Fig. 4a, Extended Data Figure 1c,e). This re-entraining stimulus was repeated for three consecutive months (Fig. 4a).

**Figure 4:**
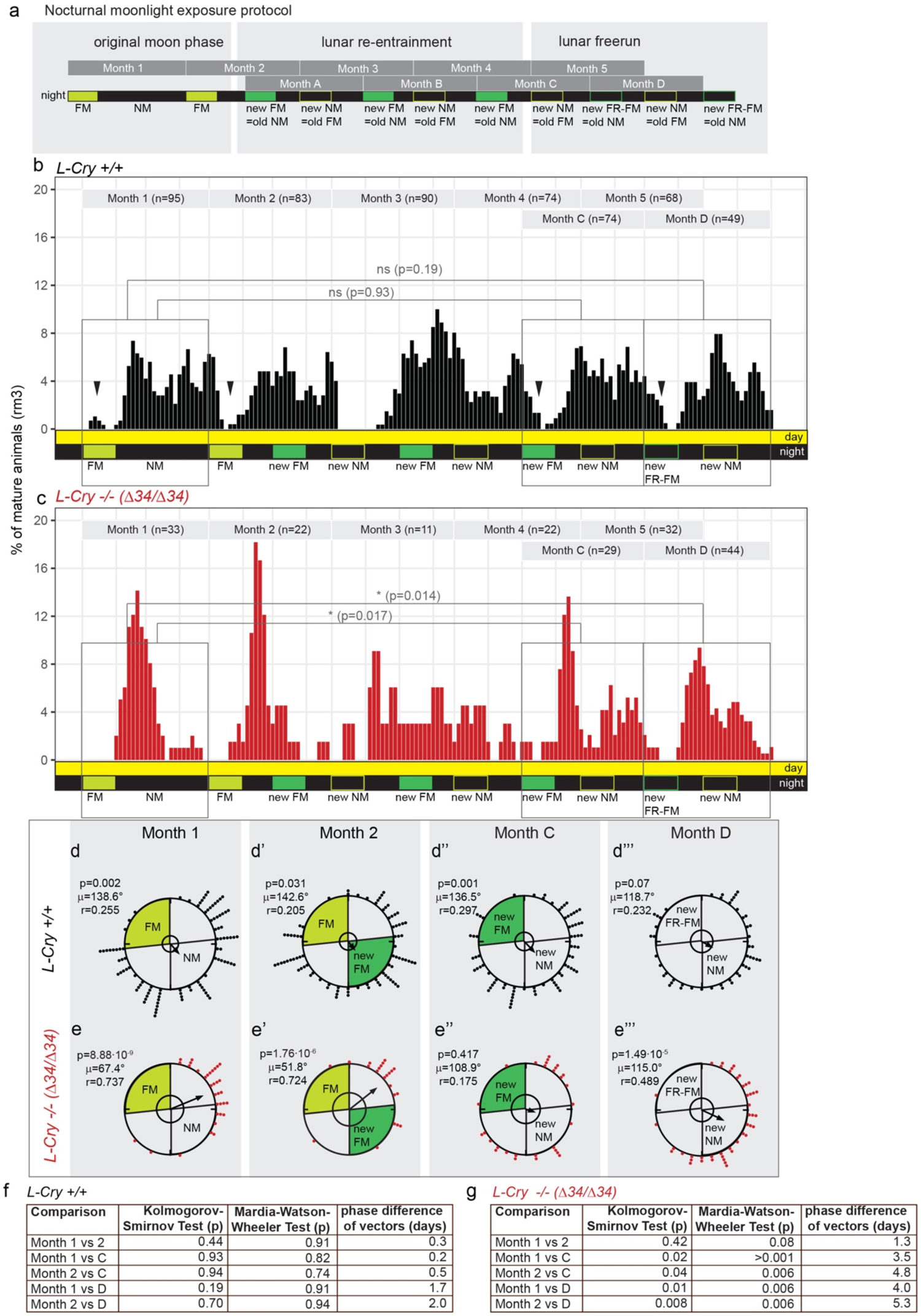
*l-cry* has a minor contribution as entraining photoreceptor to circalunar clock entrainment. **(a)** Nocturnal moonlight exposure protocol of lunar phase shift with 8 nights of naturalistic moonlight (dark green). **(b,c)** Number of mature animals (percent per month, rolling mean with a window of 3 days) of *l-cry* wildtype (b) and mutant (c) animals. p-values: Kolomogorov-Smirnov tests. Black arrowheads indicate spawning-free intervals of the wildtype, which shifted to the position of the new FM (under free-running conditions: FR-FM). **(d,e)** Data as in (b,c) plotted as circular data. 360° correspond to 30 days of the lunar month. The arrow represents the mean vector characterized by the direction angle µ and r. r (length of µ) indicates phase coherence (measure of population synchrony). p-values are results of Rayleigh Tests: Significance indicates non-random distribution of data points. The inner circle represents the Rayleigh critical value (p=0.05). **(f,g)** Results of multisample statistics on spawning data shown in (a-e). Phase differences in days can be calculated from the angle between the two mean vectors (i.e. 12°= 1 day).

The resulting spawning distribution was analysed for the efficacy of the naturalistic moonlight to phase-shift the circalunar oscillator. In order to test if the animals had shifted their spawning to the new phase, we again compared the spawning pattern before the exposure to the new fullmoon stimulus (months with numbers: data distribution analyzed relative to the initial/old FM, see scheme Fig. 4a) to the spawning pattern after the exposure to the new fullmoon stimulus (months with letters: data distribution analyzed relative to the new FM, Fig. 4a). The more similar the data distributions of month 1 is to the months C,D, the more the population was shifted to the new phase.

Wildtype animals completely shifted their spawning pattern to the naturalistic moonlight stimulus, as supported by statistical analyses: When comparing the months 1 and 2 (relative to the old FM before the shift) to the months C and D (relative to the new FM after the shift), both the Kolmogorov-Smirnov test (Fig.4b: grey rectangles, 4f) and the Mardia-Watson-Wheeler test of the same data were non-significant (Fig.4f), indicative of the population shifting to the new phase. Consistently, the direction angle (µ) of the mean vectors before and after the shift was highly similar, resulting in a phase difference of only 0.2 days between months 1 and C and 0.5 days between month 2 and month C (Fig. 4f, for details see methods).

Of note, wildtype worms would eventually reach the high spawning precision found under naturalistic moonlight only after several more months based on independent experiments (Fig. 2j,k). When we analyzed the spawning distribution of *l-cry* mutants in the same way as the wildtypes, we found that the data distribution exhibited significant differences in the linear Kolmogorov-Smirnov test when comparing months 1 and 2 before the shift to the months C and D after the shift (Fig. 4c: grey rectangles, Fig. 4g); as well as in the phase distribution in the circular analyses when comparing the months before the shift (months 1 and 2) with the last months of the shift (months C,D) (Fig. 4e,e’ versus e’’,e’’’,g). The populations also exhibited a noticeable phase difference of ≥3.5 days (Fig. 4g).

Based on the statistical significant difference in the re-entrainment of *l-cry^-/-^*, but not wild-type populations under a naturalistic sun- and moonlight regime, we conclude that L-Cry also contributes to circalunar entrainment as a photoreceptor. However, as these differences are rather minor, compared to the much stronger differences seen under artificial light regime, we conclude that its major role is the light gate keeping function.

In an independent study that focused on the impact of moonlight on daily timing, we identified r-Opsin1 as a lunar light receptor that mediates moonlight effects on the worms’ ∼24hr clock ^32^. We tested if *r-opsin1* is similarly important for mediating the moonlight effects on the monthly oscillator of the worm, analyzed here. This is not the case. *r-opsin1^-/-^* animals re-entrain as well as wildtype worms under naturalistic light conditions (Extended Data Figure 4). This adds to and is also consistent with our above observation that the spawning distribution is un-altered between *r-opsin1^-/-^* and wildtype animals under artificial light conditions (Extended Data Fig.3a,b,e,f). This finding also further enforces the notion that monthly and daily oscillators use distinct mechanisms, but both require L-Cry as light interpreter.

### L-Cry discriminates between naturalistic sun- and moonlight by forming differently photoreduced states

Given that the phenotype of *l-cry^-/-^* animals suggests a role of L-Cry as light gate keeper, i.e. only allowing the ‘right’ light to most efficiently impact on the circalunar oscillator, we next investigated how this could function on the biochemical and cell biological level.

While we have previously shown that *Pdu*-L-Cry is degraded upon light exposure in S2 cell culture ^15^, it has remained unclear if L-Cry has the spectral properties and sensitivity to sense moonlight and whether this would differ from sunlight sensation. To test this, we purified full-length L-Cry from insect cells (Extended Data Fig.5a-c). Multiangle light scattering (SEC-MALS) analyses of purified dark-state L-Cry revealed a molecular weight of 133 kDa, consistent with the predicted molecular homodimer weight of 135 kDa (Fig. 5a). Purified L-Cry binds Flavin Adenine Dinucleotide (FAD) as its chromophore (Extended Data Fig.5d,e). We then used UV/Vis absorption spectroscopy to analyze the FAD photoreaction of L-Cry. The absorption spectrum of dark-state L-Cry showed maxima at 450nm and 475nm, consistent with the presence of oxidized FAD (Extended Data Fig.5f, black line). As basic starting point to analyze its photocycle, L-Cry was photoreduced with a 445nm emitting strong blue light LED (Extended Data Fig.1d) for 110s ^33^. The light-activated spectrum showed that blue-light irradiation of L-Cry leads to the complete conversion of FAD_ox_ into an anionic FAD radical (FAD^o-^) with characteristic FAD^o-^ absorption maxima at 370 nm and 404 nm and reduced absorbance at 450 nm (Extended Data Fig.5f, blue spectrum, black arrows). In darkness, L-Cry reverted back to the dark state with time constants of 2 min (18°C), 4 min (6⁰C) and 4.7 min (ice) (Extended Data Fig.5g-k).

**Figure 5:**
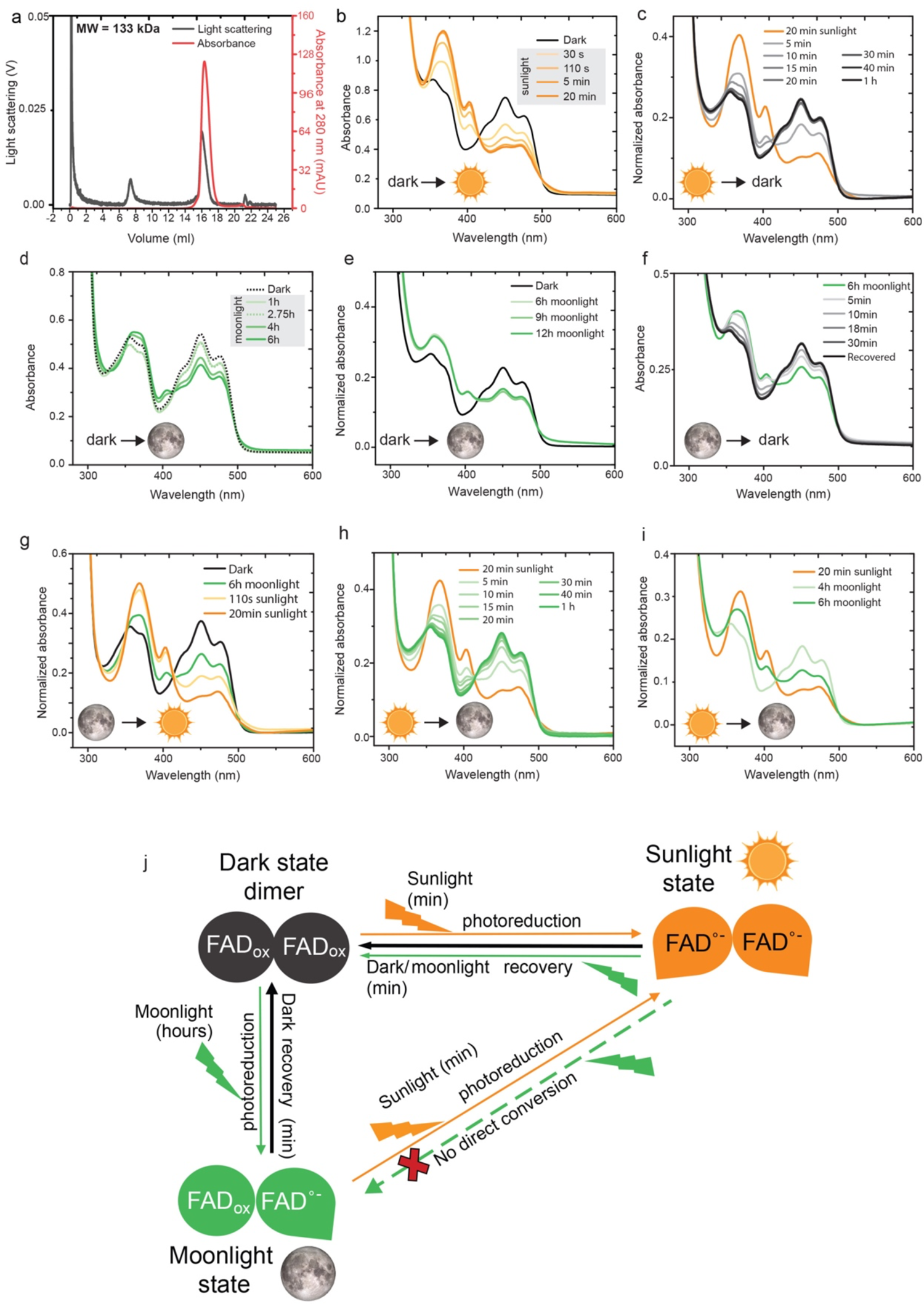
L-Cry forms differently photoreduced sunlight- and moonlight states. **(a)** Multi-Angle Light Scattering (MALS) analyses of dark-state L-Cry supports L-Cry homodimer formation (theoretical MW 135 kDa). **(b)** Absorption spectrum of L-Cry in darkness (black) and after sunlight exposure (orange). Additional timepoints: Extended Data Fig. 6a. **(c)** Dark recovery of L-Cry after 20min sunlight on ice. Absorbance at 450nm in Extended Data Fig.6b. **(d,e)** Absorption spectra of L-Cry after exposure to naturalistic moonlight for different durations. **(f)** Full spectra of dark recovery after 6h moonlight. Absorbance at 450nm: Extended Data Fig. 6d. **(g)** Absorption spectrum of L-Cry after 6h of moonlight followed by 20min of sunlight. **(h)** Absorption spectrum of L-Cry after 20min sunlight followed by moonlight first results in dark-state recovery. Absorbance at 450nm: Extended Data Fig. 6e. **(i)** Absorption spectrum of L-Cry after 20min sunlight followed by 4h and 6h moonlight builds up the moonlight state. **(j)** Schematic model of *Pdu* L-Cry responses to sun- and moonlight. MALS data (a) and SEC (Extended Data Fig.5a) suggest that L-Cry forms homodimers. Also see discussion and Extended Data Fig. 6i.

We then investigated the response of L-Cry to ecologically relevant light, i.e. sun- and moonlight using naturalistic sun- and moonlight devices we designed based on light measurements at the natural habitat of *Platynereis dumerilii* ^26^ (Extended Data Fig.1a,c,e). Upon naturalistic sunlight illumination, FAD was photoreduced to FAD°^-^, but with slower kinetics than under the blue light source, likely due to the intensity differences between the two lights (Extended Data Fig.1c-e).

While blue-light illumination led to a complete photoreduction within 110s (Extended Data Fig.5f), sunlight-induced photoreduction to FAD°^-^ was completed after 20 min (Fig. 5b) and did not further increase upon continued illumation for up to 2h (Extended Data Fig.6a). Dark recovery kinetics had time constants of 3.2min (18°C) and 5min (ice) (Fig.1c, Extended Data Fig.6b,c).

As the absorbance spectrum of L-Cry overlaps with that of moonlight at the *Platynereis* natural habitat (Extended Data Fig. 1a), L-Cry has the principle spectral prerequisite to sense moonlight. However, the most striking characteristic of moonlight is its very low intensity (1.79 x 10^10^ photons/cm^2^/s at −5m, Extended Data Fig. 1a,e). To test if *Pdu-*L-Cry is sensitive enough for moonlight, we illuminated purified L-Cry with our custom-built naturalistic moonlight, closely resembling full moon light intensity and spectrum at the *Platynereis* natural habitat (Extended Data Fig. 1a,c,e). Naturalistic moonlight exposure up to 2.75 hours did not markedly photoreduce FAD, notably there was no difference between 1 hours and 2.75 hours (Fig. 5d). However, further continuous naturalistic moonlight illumination of 4h and longer resulted in significant changes (Fig. 5d), whereby the spectrum transitioned towards the light activated state of FAD^o-^ (note peak changes at 404nm and at 450nm). This photoreduction progressed further until 6 h naturalistic moonlight exposure (Fig. 5d). No additional photoreduction could be observed after 9 h and 12 h of naturalistic moonlight exposure (Fig.5e), indicating a distinct state induced by naturalistic moonlight that reaches its maximum after ∼6hrs, when about half of the L-Cry molecules are photoreduced. This time of ∼6hrs is remarkably consistent with classical work showing that a minimum of ∼6hrs of continuous nocturnal light is important for circalunar clock entrainment, irrespective of the preceding photoperiod^5^. The dark recovery of L-Cry after 6 h moonlight exposure occurred with a time constant of 6.7 min at 18⁰C (Fig. 5f, Extended Data Fig.6d). Given that both sunlight and moonlight cause FAD photoreduction, but with different kinetics and different final FAD°^-^product/FAD_ox_ adduct ratios, we wondered how purified L-Cry would react to transitions between naturalistic sun- and moonlight (i.e. during “sunrise” and “sunset”).

Mimicking the sunrise scenario, L-Cry was first illuminated with naturalistic moonlight for 6 h followed by 20 min of sunlight exposure. This resulted in an immediate enrichment of the FAD°^-^ state (Fig. 5g). Hence, naturalistic sunlight immediately photoreduces remaining oxidized flavin molecules, that are characteristic of moonlight activated L-Cry, to FAD°^-^, to reach a distinct fully reduced sunlight state.

In contrast, when we next mimicked the day-night transition (“sunset”) by first photoreducing with naturalistic sunlight (or strong blue light) and subsequently exposed L-Cry to moonlight, L-Cry first returned to its full dark state within about 30 min (naturalistic sunlight: τ=7min (ice): Fig.5h, Extended Data Fig.6e; blue light: τ=9 min (ice): Extended Data Fig.6f-h), despite the continuous naturalistic moonlight illumination. Prolonged moonlight illumination then led to the conversion of dark-state L-Cry to the “moonlight state” (Fig. 5i, Extended Data Fig. 6f). Hence, fully photoreduced “sunlight-state” L-Cry first has to return into the dark state before entering the “moonlight-state” characterized by the stable presence of the partial FAD°^-^product/FAD_ox_ adduct. In contrast to “sunlight-state” L-Cry, “moonlight-state” L-Cry does not return to the oxidized (“dark”) state under naturalistic moonlight, i.e. moonlight maintains the “moonlight-state”, but not the “sunlight-state”. Taken together, these results indicate the existance of kinetically and structurally distinct “sunlight” and “moonlight” states of L-Cry (Fig.5j, Extended Data Fig.6i).

### Naturalistic sun- and moonlight differently affect L-Cry subcellular localization

In order to further investigate the response of L-Cry to naturalistic sun- and moonlight, we conducted Western blots and immunohistochemistry at different lunar and daily timepoints (Figs. 6a-a’’). For the analyses of total protein levels via Western blots, we compared equal lengths of sun-versus moonlight illumination versus darkness, each having 8hrs duration during their naturally occurring time (Fig.6a-a’’). L-Cry levels after 8h of naturalistic sunlight (day before full moon = FM-1, diel time: zeitgeber time 8 = zt8, see schemes in Figs. 6a,a’) were significantly reduced compared to 8h under darkness at the same moon phase (FM-1, zt 0-10mins, Figs. 6b,c), in line with (canonical) L-Cry degradation in response to naturalistic sunlight.

**Figure 6:**
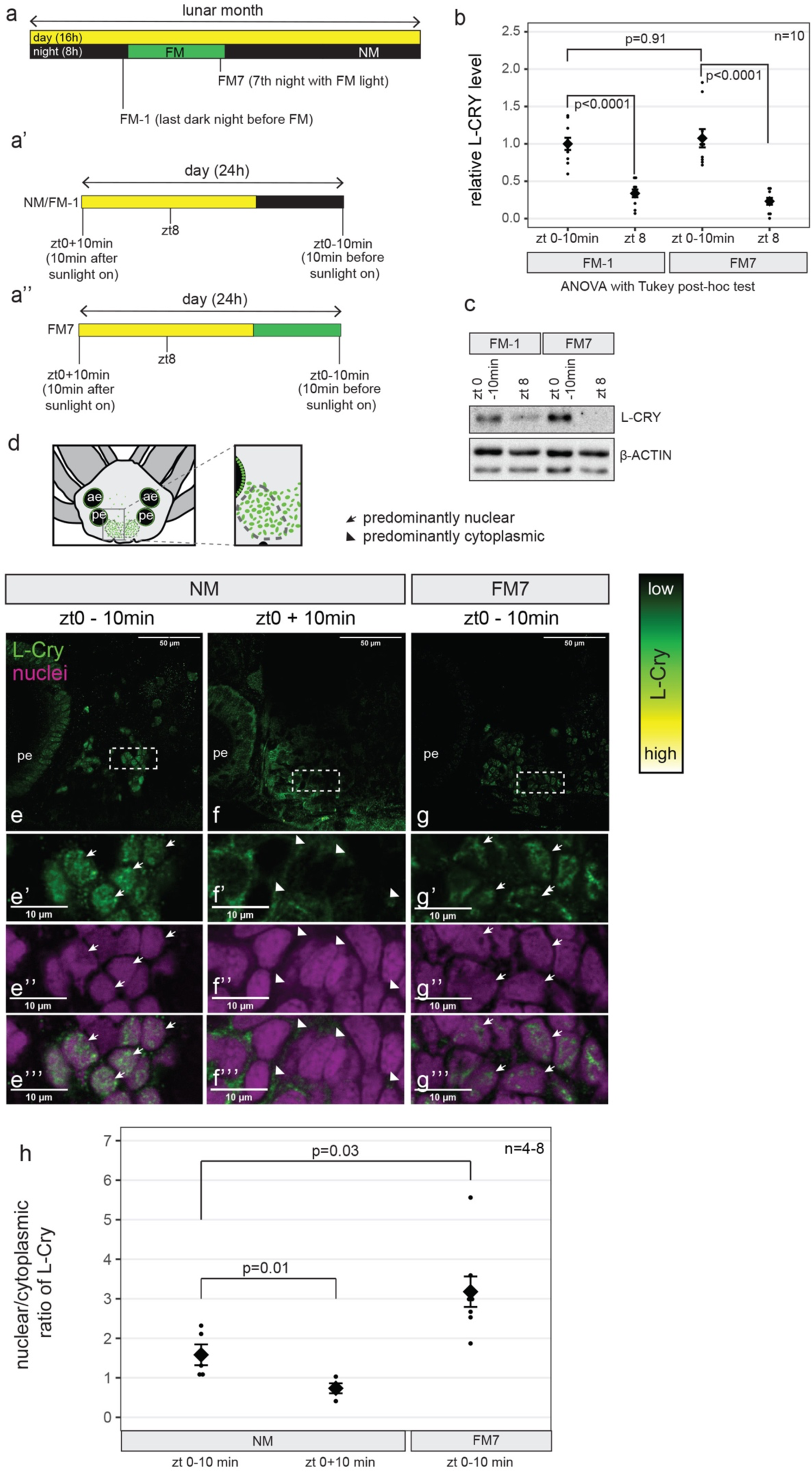
Naturalistic moon- and sunlight impact differently on L-Cry localization and levels. **(a,a’,a’’)** Scheme of sampling timepoints. 16hrs day (light) and 8 hrs night (dark or moonlight) per 24hrs, with 8 nights of moonlight per month. NM and FM-1 have the same light regime, but are named differently for accuracy as they refer to different days relative to the lunar month. **(b)** relative L-Cry levels at indicated timepoints, as determined by Western Blot. Individual data points as well as mean ± SEM are shown. **(c)** Representative Western Blot used for quantification in (b), see Extended Data Fig 7 for all other. **(d)** *P.dumerilii* head scheme. Dashed ovals designate the oval-shaped posterior domains between the posterior eyes. Green dots: L-Cry+ cells. ae, anterior eye; pe, posterior eye. **(e-g)** Confocal single layer (1.28µm) images of worm heads stained with anti-L-Cry antibody (green) and HOECHST (magenta: nuclei). White rectangles: areas of the zoom-ins presented below. **(e’-g’’’)** zoomed pictures of the areas depicted in e-g. Arrows: predominant nuclear L-Cry, arrowheads: predominant cytoplasmic L-Cry. Scale bars: 10um. Overview images with nuclear stain: Extended Data Fig 8 a-c. **(h)** quantification of subcellular localization of L-Cry as nuclear/cytoplasmic ratio at indicated timepoints. Individual data points as well as mean ± SEM are shown. p-values: two-tailed t-test. For quantification as categorical data, see Extended Data Fig 8a’-f.

In contrast to sunlight, exposure to an equal length (8hrs) of naturalistic moonlight did not cause a reduction in L-Cry levels compared to an equivalent time (8hrs) in darkness (FM-1, zt0-10min versus FM7, zt0-10min: Figs.6b,c, Extended Data Fig.7). Thus, any potential moonlight signalling via L-Cry occurs via a mechanism independent of L-Cry degradation.

We next examined the spatial distribution of L-Cry in worm heads (scheme Fig. 6d) at lunar and diel timepoints (Fig. 6a-a’’). After 8hrs of a dark night (i.e. NM, zt0-10min), L-Cry is found predominantly in the nucleus of individual cells, (Fig. 6e-e’’’, quantification as numerical data, i.e. nuclear/cytoplasmic ratio: Fig. 6h, for quantification as categorical data ^34^: Extended Data Fig. 8a’-c’’, d-f). Given that an equivalent time of 8hrs of sunlight exposure results in strong degradation of L-Cry and hence loss of staining signal (see Western blots above), we analyzed L-Cry’s localization after a short exposure. Already after 10mins of exposure to naturalistic sunlight (NM zt0+10mins, Fig. 6a,a’), the L-Cry nuclear localization strongly diminished, becoming predominantly cytoplasmic (Fig. 6f-f’’’, numerical quantification Fig. 6h, categorical quantification Extended Data Fig. 8a’-c’’, d-f). This suggests that naturalistic sunlight causes a shift of the protein to the cytoplasm, followed by degradation. (NM and FM-1 are identical in their illumination regime.)

Given the degradation of L-Cry by naturalistic sunlight, we next asked the question if L-Cry is present at night timepoints, allowing for sufficient exposure to naturalistic moonlight to reach the moonlight state. We tested two diel timepoints of the first night lit by the naturalistic moonlight for circalunar entrainment (FM1): at zt16 (just after the naturalistic sunlight is off and moonlight is on) and at zt20 (after 4hrs of naturalistic moonlight exposure) (Extended Data Fig. 9a,a’). We observe that low levels of L-Cry can already be detected at FM1 zt16 (Extended Data Fig. 9b-b’’’), and increase within the next hours (see FM1 zt20, Extended Data Fig. 9c-c’’’), with a predominantly nuclear L-Cry localization. At this timepoint still 4hours of moonlight illumination remain for the protein to biochemically reach the full moonlight state (ZT20 to ZT24). Based on these data we conclude that within the organism and under natural conditions (with the moon illuminating at least 8h of the night under full moon conditions even during summer photoperiods), L-Cry has sufficient time to reach its moonlight state (by changing from sunlight to dark to moonlight state and/or by *de novo* synthesis of dark adapted L-Cry that reaches the moonlight state within 4hrs- see biochemical kinetics, Fig.5d-j, Extended Data Figures 6f,g).

Upon further naturalistic moonlight exposure for seven continuous nights (FM7, zt0-10min) L-Cry remained clearly nuclear (Fig. 6g-g’’’, numerical quantification Fig. 6h, categorical quantification: Extended Data Fig. 8f). Thus, the sunlight and moonlight-states of L-Cry correlate with distinct subcellular distribution patterns. In fact, we observed that L-Cry at FM7, zt0-10min is even more nuclear restricted than at zt0-10min under NM, both in the numerical analysis of the nuclear/cytoplasmic ratio (Fig. 3h), as well as in the blind categorical scoring (Extended Data Fig 8f). This suggests that also the dark and moonlight states of L-Cry have distinct subcellular distribution patterns. Complementing the spawning analyses on genetically mutated animals, these findings show that moonlight and sunlight impact differentially on L-Cry quantity and localization.

This allows us to put forward a model, in which L-Cry directly via its biochemical states and connected cellular signalling properties is able to discriminate between (naturalistic) sun- and moonlight and to function as a gate-keeper for potentially entraining light stimuli for the circalunar oscillator (Fig. 7b). But why would it be required to do this in nature? As we expand in more detail in the discussion, we speculate that this is necessary to entrain to a specific moon phase, which is the full moon phase for *Platynereis*. This moon phase is specifically characterized by the long duration of detectable moonlight, i.e. moonlight during the entire night ^21^ (Fig.7a). Interestingly, this matches the biochemical kinetics of at least 6hours of light exposure to acquire L-Cry’s biochemical moonlight state. However in nature, where the setting of the full and waning moons is immediately followed by sunrise (i.e. no darkness window, Fig.7a, ^21^), measuring the duration of light exposure alone would not allow the worms to detect a specific moonphase (Fig.7a). Thus, under the natural conditions of waning/waxing moonphases and sunrise/sunsets, being able to detect the switch from moonlight to sunlight is essential to determine the end of the moonlight phase and thus to discriminate between full moon and waning moon phases (Fig.7a). Furthermore, L-Cry’s gate-keeping mechanism likely also makes the entrainment system more stable against irregular illumination as it could arise from thunderstorms.

**Figure 7.**
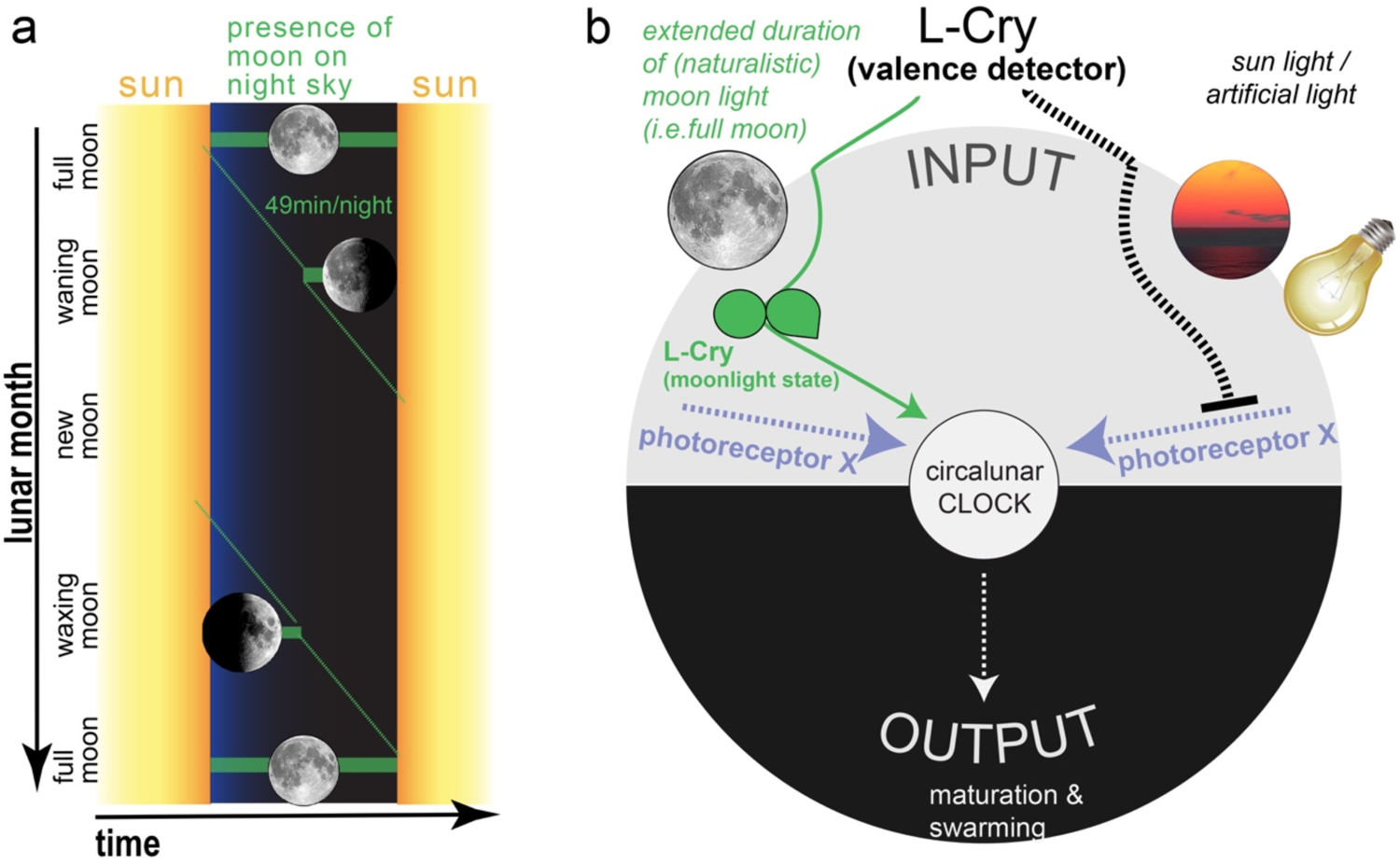
The entrainment of the monthly oscillator requires the detection of a specific moonphase. **a)** Schematic representation of the presence of the moon on the sky depending on moon phase. As a full cycle of the moon around the earth takes on average 24.8hrs, the presence of the moon relative to the sun shifts every night by ∼49mins, indicated by the green diagonal. Worms need to specifically detect the full moon phase for circalunar oscillator entrainment, which requires that they can realize when a specific light (moonlight) starts and ends ^21^. **b)** Scheme of L-Cry’s function as moonlight duration and valence detector for circalunar clock entrainment. L-Cry’s biochemical property of only reaching the full moonlight state after extended periods of (naturalistic) moonlight illumination allow for a discrimination of moonlight duration. As moon phases are characterized by the duration of the moon on the night sky, moonlight exposure duration translates into moon phase detection. L-Cry also discriminates the valence of light, strongly favoring (naturalistic) moonlight to entrain the circalunar clock.

## Discussion

Our work delivers the first molecular entry point into the mechanisms underlying a moonlight-entrained monthly oscillator. It also provides the new concept that a light receptor does not just sense light, but by its intensity and duration can give light a valence that is relevant to discriminate between different naturally existing light sources (Fig. 7a,b). While we see the most apparent behavioral/physiological phenotype of the *l-cry-/-* worms under artificial lab light conditions, these conditions are nevertheless highly informative about *l-cry’s* role as light valence detector. As briefly mentioned above, we interpret that this valence detection is under natural conditions necessary in order for the worms to synchronize to a specific moonphase: full moon. Full moon has the specific property that it is the moonphase during which the moon illuminates the entire night from sunset to sunrise (Fig.7a). In order for the organism to ‘know’ this specific moonphase under natural conditions with waning and waxing moons as well as sunrise/sunsets (Fig.7a), it needs to determine the duration not just of illumination, but specifically of the dim light illumination. This discrimination is made by L-Cry. Under the lab artificial light conditions, the moonlight stimulus is much more intense and misses signals of the waning/waxing moon phases. We hypothesize that under these artificial situations, the circalunar clock is still somewhat entrained, because there is no other entrainment stimulus it otherwise can entrain to. However, L-Cry signals that it is not really the “right” nocturnal light, which results in the observable, rather low population synchronization. If L-Cry is not present (as in the *l-cry-/-* worms), the nocturnal artificial light signal of the lab condition fully impacts on the circalunar clock. As it is (artificially) highly precise without possibly confusing waning and waxing moon signals, the entrainment results in the observed higher synchrony of the *l-cry-/-* population. If the nocturnal light signal mimics more closely the naturalistic full moon light, L-Cry permits its full impact on the circalunar oscillator, which results in the observed high population synchrony of wildtype worms under naturalistic lab light conditions. Furthermore, L-Cry’s function as a light valence detector also likely makes the entrainment system more stable against natural acute light disturbances, such as lightning.

At present, we can only speculate, how L-Cry can exert its valence function. We provide biochemical evidence that at dim light levels, corresponding to moonlight in nature, L-Cry can accumulate photons over time. L-Cry’s photoreduction response to this accumulation is in its duration markedly different from its rapid, well-established response to strong light and non-linear, suggesting that a different “moonlight” signalling state might exist. Consistent with different L-Cry biochemical states under dim and strong light, L-Cry under naturalistic moonlight is not following the conventional cytoplasmic degradation pathway, but localizes at higher levels to the nucleus. This suggests that different cellular compartments convey the different light messages to different downstream pathways.

Upon exposure to naturalistic daylight, L-Cry rapidly moves to the cytoplasm, where its protein levels become reduced, fully consistent with our previous data in S2 cells ^15^. In *Drosophila melanogaster* dCry is degraded via its light-induced interaction with the circadian clock protein Timeless and subsequent JETLAG (JET) ubiquitin-ligase–mediated proteolytic degradation ^35^. Given that both Timeless and Jetlag exist in *Platynereis* (*Pdu-*Tim: ^15^, *Pdu-*Jet: reciprocal best blast hit in ESTs), it is possible that a similar mechanism exists in *Platynereis*. It is however noteworthy that *timeless* transcript levels are affected by room light in the bristle worm ^15^, indicative of additional levels of regulatory complexity.

In contrast to the suggested mechanism about L-Cry’s fate in the cytoplasm based on existing knowledge from its *Drosophila* ortholog, the signalling downstream of nuclear L-Cry is at present completely enigmatic. It is clear from our data that neither darkness nor naturalistic moonlight causes degradation of nuclear L-Cry. This might indicate that ratios of cytoplasmic versus nuclear L-Cry could be important for circalunar clock moonlight entrainment. To complicate matters even further- it is possible that different spatial expression domains, such as eye versus brain, need to be considered separately in their responses to extended naturalistic moonlight and subsequent downstream signalling cascades.

In its role as a light-signal gate keeper, only the accumulation of L-Cry molecules in the nuclear moonlight signalling state following prolonged moonlight exposure during full-moon, would enable lunar entrainment via an additional photoreceptor “X”, which by itself is not able to discriminate between the correct full-moon signal and other “wrong” signals, such as sunlight or (in our lab experiment) the artificial/non-naturalistic nocturnal light source (Fig. 7b).

When L-Cry is photoreduced by light other than (naturalistic) moonlight, the light signalling of photoreceptor X towards the circalunar oscillator is inhibited (Fig.7b). *L-cry* mutant worms lack this inhibitory mechanism, resulting in the observed (unnatural) synchronisation to artificial moon light. On the other hand, the somewhat better response of wildtype worms to naturalistic moonlight under re-entraining conditions indicates that the accumulation of moon-light state L-Cry not only releases the inhibition, but might enhance the activity of the yet to be identified photoreceptor X or provide additional light signalling by itself to the monthly oscillator.

Connected to the question of the transmission of the moonlight signal to the circalunar oscillator is also a better understanding of L-Cry’s “moonlight state”. Is this state “just” a partial photoreduction of the state reached upon artificial sunlight exposure or perhaps also a conformationally different state with distinct formation and decay kinetics? And what is the role of the L-Cry dimers? An intriguing observation is, that in presence of moonlight the moonlight state can be stably maintained over several hours, whereas the sunlight state completely reverts to the fully-oxidized dark-state within minutes without accumulating the moonlight state while transitioning through partial photoreduction (Fig.5j). These different responses to moonlight illumination suggest that the moonlight- and sunlight states are conformationally and kinetically not equivalent (Extended Data Figure 6i). Based on its sequence homology and the similarity of its FAD photoreaction to *Drosophila* CRY (dCry), it is conceivable that L-Cry also displaces the regulatory C-terminal tail in the photoreduced state as observed for dCry ^33, 36^. However, as dCry is monomeric, L-Cry homodimer formation may impact these conformational changes, and these may further vary depending on whether moonlight or sunlight operates on the initial dark-state L-Cry homodimer. We propose, that partial FAD photoreduction in the moonlight state could be related to the formation of asymmetric L-Cry dimers, where one monomer retains oxidized FAD, while in the second monomer FAD is photoreduced to FAD°^-^ (Fig. 5j). This requires, that the flavins in the two L-Cry monomers have different redox potentials, likely resulting from different chemical environments due to conformational differences between the monomers (Extended Data Figure 6i). Hence different amounts of energy (photon numbers) would be needed to photoreduce the flavins in the two L-CRY monomers. Moonlight, due to its very low intensity can only induce the lower energy transition, resulting in the partially photoreduced moonlight state. In presence of intense sunlight, however, the larger energy barrier to photoreduce the second flavin can also be overcome. Certainly, more extensive mechanistic studies are required to further support our model. However, this model is consistent with all our current *in vitro* data, and moreover, it plausibly illustrates how the very different intensities of moon- and sunlight can lead to the formation of conformationally distinct dark state (new moon), moonlight state (full moon) and sunlight state L-Cry proteins. Thereby L-Cry could translate different light qualities into different cellular signaling events, e.g. by changing L-Cry’s subcellular localizations and cellular degradation rates (Fig. 6), to ultimately affect moonlight dependent physiology (Fig. 2-4).

Finally, an evolutionary consideration: Monthly synchronization by the moon has been documented for a wide range of organisms-including brown and green algae, corals, crustaceans, worms, but also vertebrates (reviewed in ^6^). Furthermore, recent reports also provide increasing evidence that the lunar cycle influences human behaviour (reviewed in ^21, 37^). Are the lunar effects mediated by conserved or different mechanisms?

When considering monthly oscillators with period lengths in the range of weeks, our implication of L-Cry as a light receptor in the circalunar entrainment pathway at first glance rather suggests that such monthly oscillator might not be conserved, given that direct L-Cry orthologs are not present in all the groups that are affected by the lunar cycle ^38^. However, taking further aspects into account, such a conclusion might be too quick. Could other members of the Cry/photolyase family take over similar functions? Furthermore, our entrainment data suggest the presence of additional moonlight entrainment photoreceptors, which might be conserved. Last, but not least the molecular mechanisms underlying the circalunar oscillator also await identification, and it is possible that conservation exists on this level. Examples are known from circadian biology and it will now require further work to reach a similar level of understanding for moon-controlled monthly rhythms and clocks.

## Materials and Methods

### Natural light measurements

Under water measurements of natural light at the habitat of *Platynereis dumerilii* were acquired using a RAMSES-ACC-VIS hyperspectral radiometer (TriOS GmbH) for UV to IR spectral range. In coastal waters of the Island of Ischia, in the Gulf of Naples, the two radiometers were placed on sand flat at 5m depth near to *Posidonia oceanica* meadows, which are a natural habitat for *P. dumerilii.* Measurements were recorded automatically every 15min across several weeks in the winter 2011/2012. To obtain a fullmoon spectrum, measurements taken from 10pm to 1am on a clear fullmoon night on the 10.11.2011 were averaged. To subtract baseline noise from this measurement, a NM spectrum was obtained by averaging measurements between 7:15pm to 5am on a NM night on 24.11.2011, and subtracted from the FM spectrum. Resulting spectrum: Extended Data Fig. 1a. To benchmark these moonlight spectra measured under water with moonlight measured on land, we compared the underwater spectra to a publicly available full moon spectrum measured on land on 14.04.2014 in the Netherlands (Extended Data Fig. 1g, spectrum available at http://www.olino.org/blog/us/articles/2015/10/05/spectrum-of-moon-light). As expected, light with longer wavelengths was strongly reduced in the underwater measurements compared to the surface spectrum, since longer wavelengths penetrate water less efficiently. For the sunlight spectrum, measurements taken from 8am to 4pm on a sunny day on 9.11.2011 were averaged.

### Naturalistic Light Systems (NELIS devices)

To emulate naturalistic sunlight and moonlight conditions, we employed NELIS (Natural Environment Light Intensity System) (Marine Breeding Systems GmbH)^26^. The naturalistic moonlight device was composed of a combination of LEDs and an Ulbricht sphere for homogenous light mixing. For improved light distribution across the shelf, a naturalistic moonlight device was attached to each end of an acrylglass rod (two light sources and one rod per shelf). For details on the naturalistic sunlight source see ref. ^26^. Light spectra were measured using a ILT950 Spectroradiometer (International Light Technologies).

### Cloning and recombinant virus generation for L-Cry

Full length N-terminally His_6_-tagged *Platynereis dumerilii* L-Cry (1-567) was heterologously expressed in *Spodoptera frugiperda (Sf9)* insect cells using the Bac-to-Bac baculovirus expression system with the pCoofy27 expression vector. 1 * 10^6^ *Sf9* cells were transfected with recombinant bacmid DNA using Cellfectin. The first generation P0 virus was harvested 3 days after bacmid transfection. A further virus amplification step was carried out and the P1 virus stock was used for protein expression. The volume of P1 virus stock to be added for sufficient protein expression was determined by test expression.

### Protein expression and purification

*Sf9* cells were grown as suspension cultures in Sf-900 II media at 27°C, 80 RPM. 1 L of 1 * 10^6^ *Sf9* cells/ml were transfected with P1 virus stock and incubated at 27°C for 72 h. Cells were harvested by centrifugation at 7000 rpm for 20 min and stored at −80°C until purification. All purification steps were carried out in dark or dim red light conditions. Columns were wrapped with aluminum foil to avoid light-activation of L-Cry. The cell pellets were resuspended in lysis buffer (20 mM Tris pH 7.5, 150 mM NaCl, 20 mM imidazole, 5% glycerol, 5 mM β-mercaptoethanol) and lysed using a microfluidizer. The lysate was centrifuged at 27000 rpm for 45 min and the clarified supernatant incubated with nickel beads for 1 h. The nickel beads were loaded onto a batch column, washed with 50-100 mM imidazol and the L-Cry protein was eluted with 250 mM imidazole. Elution fractions containing L-Cry were concentrated, diluted with low salt buffer (50 mM Tris pH 7.5, 50 mM NaCl, 5% glycerol, 1mM DTT) and loaded onto a 5 ml Hitrap Q sepharose anion exchange column (GE Healthcare). A gradient from 0 % to 100 % high salt buffer (50 mM Tris pH 7.5, 1 M NaCl, 5% glycerol, 1mM DTT) was applied. L-Cry containing fractions were pooled, concentrated and loaded onto a HiLoad S200 16/60 size exclusion chromatography (SEC) column (buffer 25 mM Bis-Tris propane pH 8.0, 150 mM NaCl, 5% glycerol, 1 mM TCEP). Fractions containing pure L-Cry were pooled, concentrated to 10 mg/ml and snap frozen in liquid nitrogen for storage at −80°C. 2 mg of L-Cry was obtained from 10 g of pellet. The identity of the L-Cry protein was confirmed by mass spectrometry.

### Reverse-phase HPLC analyses of the chromophore content of L-Cry

Flavin Mononucleotide (FMN), Flavin Adenine Dinucleotide (FAD) and Methenyltetrahydrofolate (MTHF) were dissolved in buffer (25 mM Bis-Tris propane pH 8.0, 150 mM NaCl, 5% glycerol) and run at 1ml/min (20 °C) over a Macherey-Nagel C18 Gravity-SB (150/4/5 µm) column to separate the chromophores by reverse phase (RP) HPLC analyses. A gradient from 20-100% of methanol against water (+0.1% Trifluoroacetic acid) was used for optimal separation. To analyse the chromophore content of L-Cry, purified L-Cry was heat-denatured for 5 min at 97°C and centrifuged at 14000 RPM for 10 min at 4°C. The supernatant was subjected to RP-HPLC analysis. The chromophores were monitored by absorption at 370 nm.

### Analytical Size Exclusion Chromatography (SEC) and SEC coupled with Multiangle light scattering (SEC-MALS)

Analytical SEC of dark-state L-Cry was carried out on a S200 10/300 size exclusion column (SEC buffer 25 mM Bis-Tris propane pH 8.0, 150 mM NaCl) under red light conditions. SEC-MALS was carried out to determine the exact molecular weight and oligomeric state of purified L-Cry based on the SEC elution volume and light scattering. For SEC-MALS, purified L-Cry was loaded onto a Superose 6 10/300 size exclusion column and run at a flowrate of 0.4 ml/min in SEC buffer. MALS data were obtained from a DAWN DSP instrument (Wyatt Tech, Germany) and processed using ASTRA 4.90.07. Elution volumes and corresponding molecular weight of calibration standards: 10.3 ml – 670 kDa (Thyroglobulin), 13.67 ml – 158 kDa (γ-globulin), 15.71 ml – 44 kDa (ovalbumin), 17.42 ml – 17 kDa (myoglobin) and 20.11 ml – 1350 Da (vitamin B12).

### UV/VIS spectroscopy on L-Cry: Blue light-, sunlight- and moonlight photoreduction and dark recovery

UV/Visible absorption spectra of the purified L-Cry protein were recorded on a Tecan Spark 20M plate reader unless otherwise stated. A light-state spectrum of L-Cry with fully photoreduced FAD°^-^ was collected after illuminating dark-adapted L-Cry for 110 sec with a 450 nm blue light emitting diode (Extended Data Fig. 1 d,e; 6.21 x 10^16^ photons/cm^2^/sec at the sample). To analyze sunlight- and moonlight dependent FAD photoreduction, dark-adapted L-Cry (kept on ice) was continuously illuminated with naturalistic sunlight (Extended Data Fig. 1 c,e; 1.55 x 10^15^ photons/cm^2^/sec at the sample) or naturalistic moonlight (Extended Data Fig. 1 c,e; 9.65 x 10^10^ photons/cm^2^/sec at the sample) and UV-VIS spectra (300 – 700 nm) were collected at different time points.

Dark recovery kinetics (FAD reoxidation) of L-Cry at 18°C following illumination with blue-light (110 sec), sunlight (20 min on ice) or moonlight (6 h on ice) were measured by recording absorbance changes at 450 nm over time or by extracting 450 nm absorbance values from complete UV/VIS spectra. To measure dark recovery kinetics on ice, complete UV-VIS spectra (300 – 700 nm) were collected at different time points following 110 sec blue light- or 20 min sunlight illumination and absorbance values at 450 nm were extracted from the full spectra (sample was kept on ice and in darkness between measurements). Additionally, a temperature controlled Jasco V-550 UV-VIS spectrophotometer was used to determine dark recovery kinetics of L-Cry (after 110 sec blue-light) at 6°C based on absorbance changes at 450 nm. The time constants for dark recovery were calculated by fitting a single exponential curve to the experimental data. Spectra were analyzed using Origin (Version 7.5/10.5(trial); OriginLab Corporation, Northampton, MA, USA).

### Recovery of L-Cry dark state in presence of moonlight

To assess if moonlight can maintain the light state, L-Cry was initially illuminated with sunlight for 20 min or with blue light for 110 sec, followed by continuous moonlight illumination up to 6 hours with the sample kept on ice. Complete UV-VIS spectra (300 – 700 nm) were collected at different time points. Absorbance values at 450 nm were taken from the complete spectra obtained between 5 min and 2h 30 min moonlight exposure and used to determine the time constant for recovery of oxidized FAD after blue-light- or sunlight induced photoreduction in presence of moonlight.

### Sunlight illumination of moonlight activated L-Cry

To assess if sunlight can further increase FAD photoreduction starting from the moonlight activated state, L-CRY was first illuminated with continuous moonlight for 6 hours, followed by 20 min of sunlight illumination (on ice). Complete UV-VIS spectra from 300 – 700 nm were measured in each case.

### Worm Culture

*Platynereis dumerilii* were grown as previously described ^15, 39^. All animal work was conducted according to Austrian and European guidelines for animal research. Photoperiod 16:8 (L/D), circalunar entrainment: nocturnal light for 6-8 nights (see Figure legends for each experiment) every 29 to 30 days (centering around full moon (“inphase”) or new moon (“outphase”) in Vienna). Light spectra and intensities of Extended Data Fig. 1 were measured with a recently calibrated ILT950 spectrometer (International Light Technologies Inc Peabody, USA) and converted to photons/cm^2^/s.

### Generation and Genotyping of *l-cry* KO worms

Design and construction of TALENs targeting *l-cry* is described in ^23^. For genotyping, DNA extraction of immature and premature worms was conducted by cutting 5-10 tail segments with a scalpel and incubating them in 20µl 50mM NaOH at 95° for 20min. After adding 5µl of Tris/HCl pH 7.5, the supernatant was used as template for the PCR reaction. Mature worms were frozen as whole at − 20°C and DNA was later extracted using NucleoSpin Tissue Mini kit for DNA from cells and tissue (Macherey-Nagel). PCR was performed with OneTaq Quick-Load 2x Master Mix with Standard Buffer (New England Biolabs). PCR product was run on an agarose gel and genotype was determined on size (168bp: wildtype, 134bp: Δ34+Δ9 mutant allele, 157bp: Δ11 mutant allele).

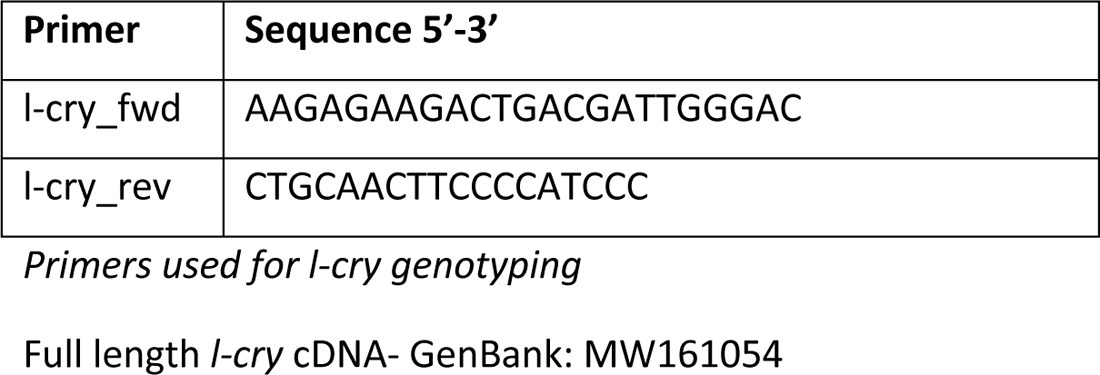

### Monoclonal Antibody Production

A peptide consisting of amino acids 52 - 290 of L-Cry protein (GenBank ID: MT656570, predicted size 25kDa) was cloned and expressed in bacteria cells. Subsequently, this peptide was purified and used for mouse immunization, thereby acting as epitope in production of a monoclonal antibody against L-Cry. Upon screening of multiple clones, two clones (4D4-3E12-E7 and 5E3-3E6-E8) were selected and used in combination. Monoclonal antibodies were produced by and purchased from the Monoclonal Antibody Facility at Max Perutz Labs (Medical University of Vienna, Department of Medical Biochemistry).

### Immunohistochemistry, microscopy and L-Cry localization determination

Worm heads were dissected with jaws and fixed in 4% PFA for 24h at 4°C. Samples were subsequently permeabilized using methanol, digested for 5 min with Proteinase K at room temperature without shaking and post-fixed with 4% PFA for 20 min at room temperature. Next, samples were washed 5 times for 5 min with 1x PTW and incubated in hybridization mixture^40^ used in *in situ* hybridization protocol, at 65° C overnight. Worm heads were washed with 50% formamide/2X SSCT - standard saline citrate containing 0.1% Tween 20® (Sigma Aldrich) (2x, 20 min), then with 2X SSCT (2x, 10 min) and with 0.2X SSCT (2x, 20 min); all washing steps at 65° C. After blocking for 90 min with 5% sheep serum (Sigma-Aldrich) at room temperature, samples were incubated in L-Cry antibodies 5E3-3E6-E8 (1:100) and 4D4-3E12-E7 (1:50) in 5% sheep serum (Sigma-Aldrich). Secondary antibody, Cy3 goat anti-mouse IgG (A10521, Thermo Fisher Scientific) was diluted 1:400 in 2.5% sheep serum (Sigma-Aldrich). Incubations were done for at least 36h at 4° C shaking and after each incubation time, samples were washed with 1x PTW three times for 15 min at room temperature and a fourth time over night at 4° C. After this, Höchst 33342 (H3570, Thermo Fisher Scientific), diluted 1:2000, was added for at least 30 min at room temperature. Samples were then washed three times for 15 min with 1x PTW at room temperature and mounted with 25 mg/ml DABCO (Roth/Lactan) in an 87% glycerol (Sigma-Aldrich) solution. All solutions were made with 1x PTW (PBS + 0.1% Tween 20®) (Sigma Aldrich). Heads were imaged on a Zeiss LSM 700 laser scanning confocal microscope using LD LCI Plan-Apochromat 25X, Plan-Apochromat 40X by CHD: T-PMT detection system and Zeiss ZEN 2012 software. Lasers: DAPI 405 nm and Cy3 555 nm.

Categorical scoring: Using Fiji/ImageJ ^41^, nuclear outlines were marked as Regions Of Interest (ROI) on the 405 nm channel images (Höchst staining). ROIs were then used for scoring of the signal localization (inside =nucleus versus outside= cytoplasm) on the 555nm channel of the same images (L-Cry).

Quantitative scoring: Using the deep learning-based image segmentation algorithm Cellpose ^42^ on the 405 nm channel images, the Hoechst-stained nuclei were identified and marked as Regions Of Interest (ROI). L-Cry signal was then determined for these nuclear ROIs using in Fiji/ImageJ ^41^. Signal intensity was determined by calculating Corrected Total Cell Fluorescence (CTCF) using the formula CTCF=Area (ROI_1)*Mean (ROI_1)-Area (ROI_1)*Mean(ROI_(background ROIs)). A sum of CTCF values of all the nuclei was subtracted from the CTCF value of the whole brain area, to obtain the corresponding value for non-nuclear, i.e. cytoplasmic signal. Finally, the ratio between nuclear and non-nuclear (cytoplasmic) signal intensity was calculated for corresponding regions of different worm heads to compare between different ZTs.

### Protein extraction and Western Blots

Per biological replicate four premature worms were anaesthetized (7.5% MgCl_2_-/H_2_0, 1:1 diluted with sea water), decapitated and heads transferred to a 1.5ml tube containing 150 µl RIPA lysis buffer (R0278 Sigma-Aldrich), 10% Triton X100 and protease inhibitor (cOmplete Tablets, EDTA-free, *EASYpack*, Roche). The tissue was homogenized by grinding using a tight fitting pestle. All steps on ice. Cell debris was pelleted by centrifugation. Protein concentration of lysates was determined using Bradford reagent (BIORAD), subjected to SDS-gel electrophoresis (10% Acrylamide) and transferred (Transferbuffer: 39mM Gylcine, 48mM Tris, 0.04% SDS, 20% MetOH) to a nitrocellulose membrane (Amersham™ Protran™ 0,45μm NC, GE Healthcare Lifescience). Quality of transfer was checked by Ponceau-S (Sigma Aldrich) staining. After 1h of blocking with 5% slim milk powder (Fixmilch Instant, Maresi) in 1xPTW (1xPBS/0.1% TWEEN 20) at room temperature, the membrane was incubated with the appropriate primary antibody diluted in 2.5% milk/PTW at 4°C overnight. [anti-L-Cry 5E3-3E6-E8 (1:100) and anti-L-Cry 4D4-3E12-E7 (1:100); anti-beta-Actin (Sigma, A-2066, 1: 20.000)]. After 3 rinses with 1xPTW the membrane was incubated with the species specific secondary antibody [anti-Mouse IgG-Peroxidase antibody, (Sigma, A4416, 1:7500); Anti-rabbit IgG-HRP-linked antibody (Cell Signaling Technology, #7074, 1:7.500] diluted in 1xPTW/1% slim milk powder for 1 hour, RT. After washing, SuperSignal™ West Femto Maximum Sensitivity Substrate kit (Thermo Fisher Scientific) was used for HRP-signal detection and finally signals were visualized by ChemiDoc Imaging System (BIORAD). Specific protein bands were quantified in “Image J” and L-Cry was normalized to beta-Actin.

### Collection and analysis of spawning data

Worm boxes were checked daily for mature worms. Worms which had metamorphosed into their sexually mature male or female form and had left their tube to perform their nuptial dance were scored as mature animals.

The recordings of mature animals in nature (collected from June 1929 to June 1930 in Naples ^29, 31^) were digitalized and all months were aligned to relative to the same moonphase and combined. For comparisons of these data with our spawning data from the lab, we aligned the first day after full moon in nature with the last day of full moon stimulus in the lab, since *Platynereis dumerilii* synchronizes its circalunar clock to the end of the full moon stimulus^5^.

For analysis, each day of the lunar month was assigned a number from 1 to 30. For linear plots, the percentage of mature worms per lunar day was then plotted as a histogram. The spawning distributions of two conditions were compared using the Kolmogorov-Smirnov Test. For the circular analysis ^43–45^ of spawning data, the lunar day of spawning was multiplied by 12 for each worm, so that the 30 lunar days regularly distributed on the 360° circle. Each dot represents one mature worm unless stated otherwise. Circular data can be described using the mean vector (displayed as an arrow), which is defined by its direction angle (µ) and its length (r). The direction angle µ is given relative to 0° (moon off). The value of length r (also called phase coherence) ranges from 0 to 1, where higher values indicate higher phase coherence (i.e synchrony). In order to test, if the observed data distribution is significantly different from random, we performed the Rayleigh’s Uniformity Test and used p<0.05 as cutoff for significance. Non-uniform distribution is consistent with lunar rhythmicity. For comparing two circular datasets (e.g. of different genotypes or different months in the phase shift experiments), we used the non-parametric Mardia-Watson-Wheeler test. Circular analysis of these data was performed using Oriana (Version 4.02, Kovach Computing Services).

### Phase-Shift Experiments

For Phase-Shift experiments, boxes with adult worms (at least 3 months old) were transferred from standard light conditions (see “worm culture”) to the naturalistic light systems (sun- and moonlight) mounted in light-tight black shelves. Number of mature worms was recorded daily and number of mature worms per day was used to calculate percentage of mature worms (one month: 100%). Data were smoothened using a rolling mean with a window size of 3 days. Data analysis was performed as described in “Collection and analysis of spawning data”.

### Whole mount *in situ* Hybridisation combined with immunohistochemistry for L-Cry

Probes were generated *de novo* using previously cloned plasmids as template. Genes of interest were amplified via PCR using Phusion Polymerase (NEB) and primers for pJET 1.2 with an overhang of Sp6 promoter. PCR product was purified following the protocol for “QIAquick PCR Purification Kit” (Qiagen). Agarose gel electrophoresis showed right amplicon sizes and single bands. For the in vitro transcription, 1µg of linearized template was used. Riboprobes were labelled with anti-Digoxigenin UTPs (Roche Diagnostics) and transcribed with Sp6 polymerase at 37°C for 4h. Probes were purified according to the “RNeasy Kit” (QIAGEN) and eluted in 40ul RNase-free water. 600-1000ng of the riboprobes were used.

Whole-mount *in situ* Hybridisation was carried out on premature worms of the RE strain, following published procedures ^40, 46^ with adjustments made to combine it with L-Cry immunohistochemistry. Worm heads were fixed in 4% PFA for 2h at RT while shaking. Proteinase K digest: 5min, during which samples were very slightly rocked. After blocking in 5% sheep serum/1X PTW, worm heads were incubated with the monoclonal L-Cry antibodies (5E3-3E6-E8 diluted 1:100 and 4D4-3E12-E7 diluted 1:50), anti-Digoxigenin-AP coupled antibody (Roche Diagnostics) and sheep serum diluted to 2.5% with 1xPTW for 36-40h at 4°C, shaking. After detection using NBT/BCIP, samples were incubated in the secondary antibody Alexa Fluor-488 goat anti-mouse IgG (Thermo Fisher Scientific), 1:400 (36-40h at 4°C, shaking), washed in 1xPTW and mounted in DABCO/Glycerol. Imaging was done using Axioplan Z2 Microscope (Carl Zeiss) with AxioCam MRc5 colour CCD camera (Carl Zeiss) and captured using ZenPro Software (Carl Zeiss). The images were edited with either ImageJ or Photoshop CC.

## Statistical Analysis

All data analysis was conducted using R 3.6.1^47^, GraphPad Prism 8.4.2, Oriana 4.02 and Microsoft Excel 2010.

## Acknowledgements

We thank the members of the Tessmar-Raible and Wolf groups for discussions. Andrej Belokurov and Margaryta Borysova for excellent worm care at the MFPL aquatic facility. We are grateful for support by the IMB Media Lab, Protein Production and Proteomics Core Facilities (mass spec instrument funded by DFG INST 247/766-1 FUGG). We would like to thank Dr Svenja Morsbach and Beate Müller from the Max Planck Institute for Polymer Research for helping with HPLC experiment and Prof Dr Elmar Jaenicke from University of Mainz for help with SEC-MALS.

## Extended Data Figures

**Extended Data Figure 1:**
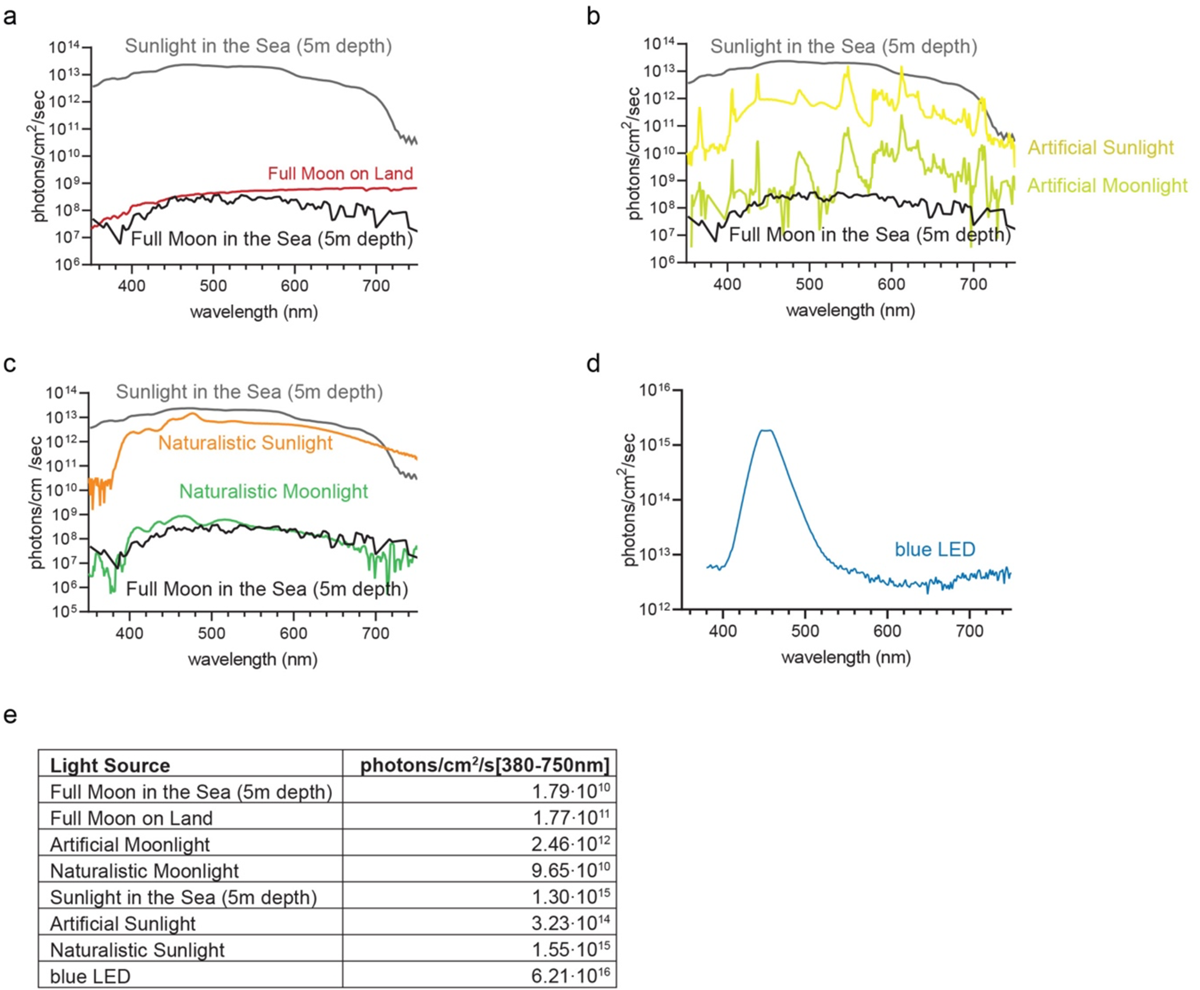
Spectra of light sources. **(a)** Light spectra in nature: sunlight (grey) and full moon in 5m depth in Ischia (black) as well as full moon light on land (red). **(b)** Spectra of the highly artificial sun-(yellow) and moonlight (light green) in the worm culture room. **(c)** Spectra of designed naturalistic sun- (orange) and moonlight (green). **(d)** Spectrum of blue light LED used for spectroscopic experiments. **(e)** number of photons/cm^2^/s of (a-d). Intensity and spectrum were always measured in the distance relevant for the experiments.

**Extended Data Figure 2:**
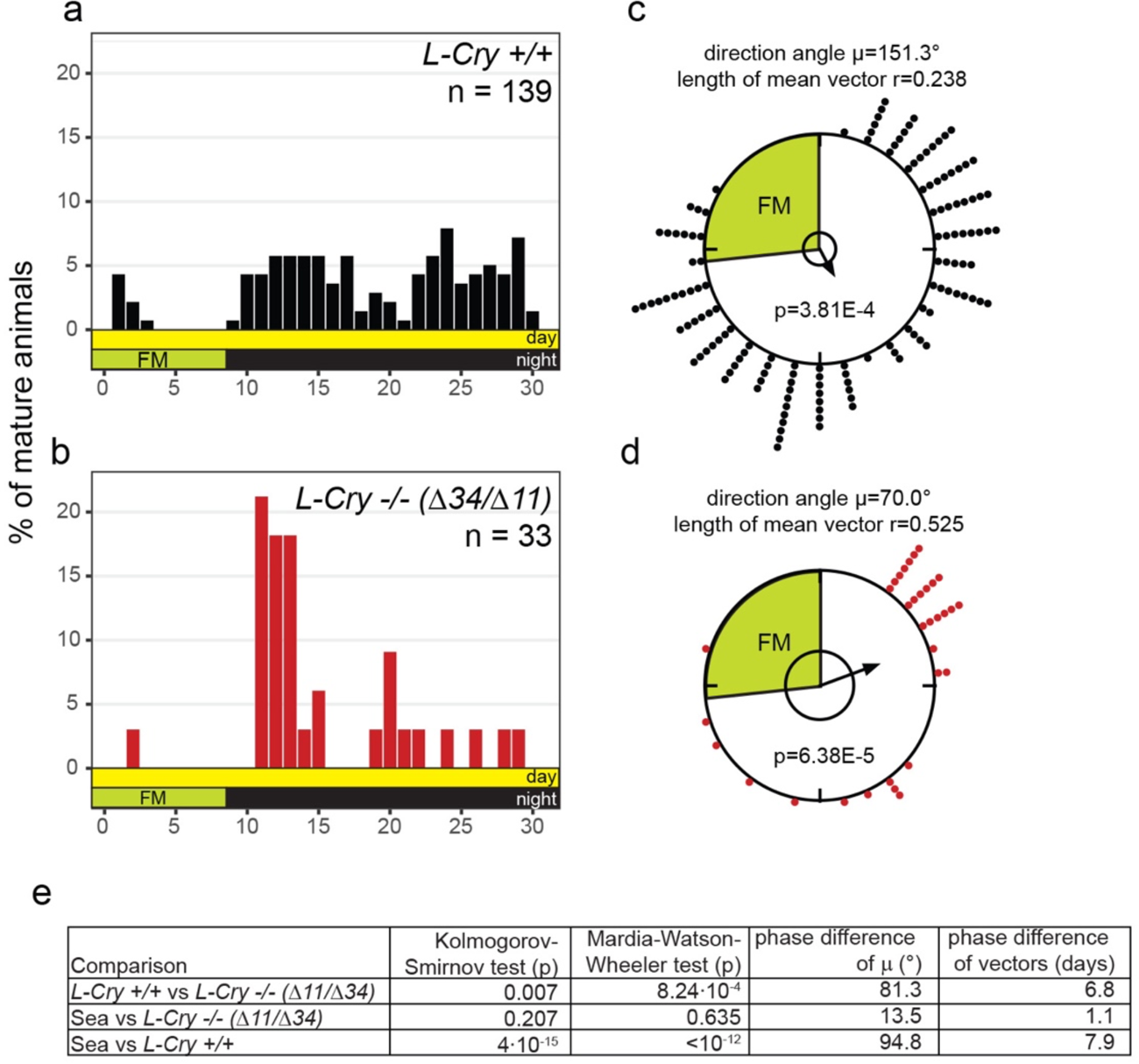
*l-cry* transheterozygous (Δ11/Δ34) mutants show increased spawning synchrony. **(a-c)** Spawning of *l-cry +/+* (a) and *l-cry -/-(Δ11/Δ34)* (b) animals over the lunar month under 8 nights of standard worm room culture full moon. **(c,d)** Same data as in (a,b) plotted as circular data. 360° correspond to 30 days of the lunar month. The arrow represents the mean vector characterized by the direction angle µ and r. r (length of µ) indicates phase coherence (measure of population synchrony). p-values inside the plots are results of Rayleigh Tests: Significance indicates non-random distribution of data points. The inner circle represents the Rayleigh critical value (p=0.05). **(e)** Results of multi-sample statistics on spawning data shown in (a-d). Phase differences in days can be calculated from the angle between the two mean vectors (i.e. 12° = 1 day).

**Extended Data Figure 3:**
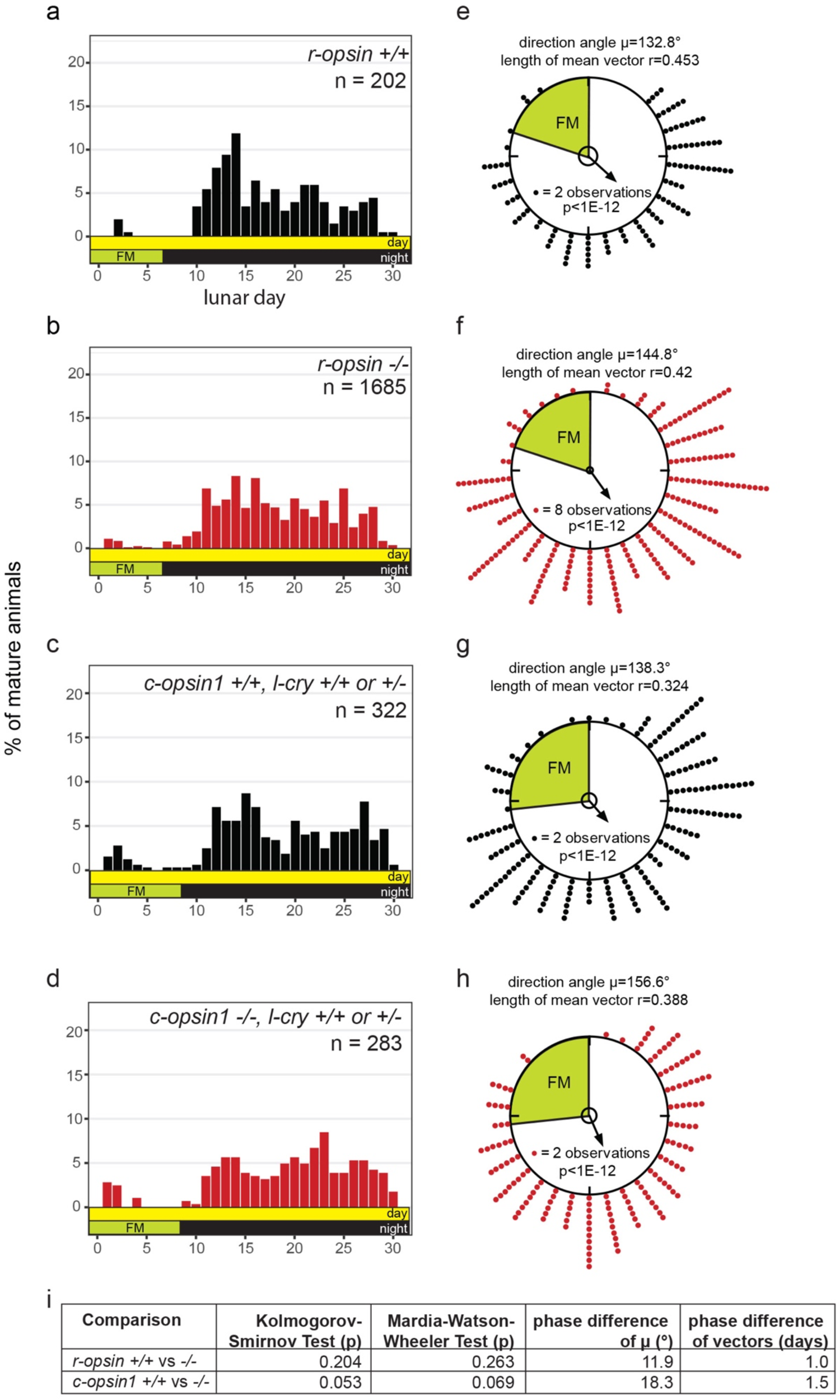
*r-opsin* and *c-opsin1* mutants show no alterations in spawning synchrony. **(a-d)** Spawning of indicated genotypes under standard laboratory conditions with 6 nights (*r-opsin*) or 8 nights (*c-opsin1*) of full moon. Animals in c and d are *l-cry +/+* or *+/-.* **(e-h)** Same data as in (a-d) plotted as circular data. 360° correspond to 30 days of the lunar month. The arrow represents the mean vector characterized by the direction angle µ and r. r (length of µ) indicates phase coherence (measure of population synchrony). p-values inside the plots are results of Rayleigh Tests: Significance indicates non-random distribution of data points. The inner circle represents the Rayleigh critical value (p=0.05). **(i)** Results of multi-sample statistics on spawning data shown in (a-h). Phase differences in days can be calculated from the angle between the two mean vectors (i.e. 12° = 1 day).

**Extended Data Figure 4:**
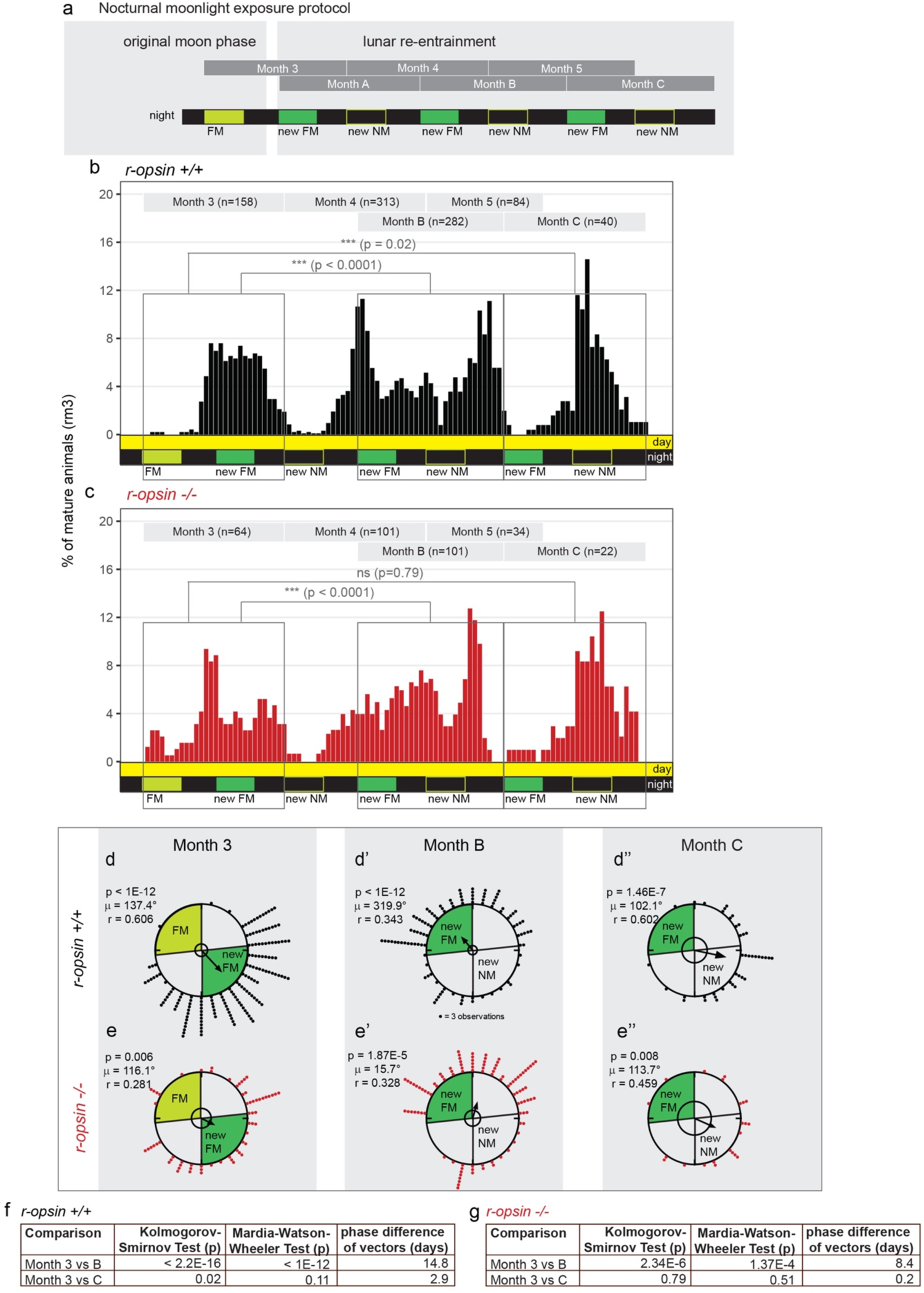
The photoreceptor molecule *r-opsin1* does not contribute to circalunar clock entrainment. **(a)** Nocturnal moonlight exposure protocol of lunar phase shift with 8 nights of naturalistic moonlight (dark green). **(b,c)** Number of mature animals (percent per month, rolling mean with a window of 3 days) of r-opsin wildtype **(b)** and mutant **(c)** animals. p-values: Kolomogorov-Smirnov tests. **(d,e)** Data as in **(b,c)** plotted as circular data. 360° correspond to 30 days of the lunar month. The arrow represents the mean vector characterized by the direction angle µ and r. r (length of µ) indicates phase coherence (measure of population synchrony). p-values are results of Rayleigh Tests: Significance indicates non-random distribution of data points. The inner circle represents the Rayleigh critical value (p=0.05). **(f,g**) Results of multisample statistics on spawning data shown in **(a-e).** Phase differences in days were calculated from the angle between the two mean vectors (i.e. 12°= 1 day).

**Extended Data Figure 5:**
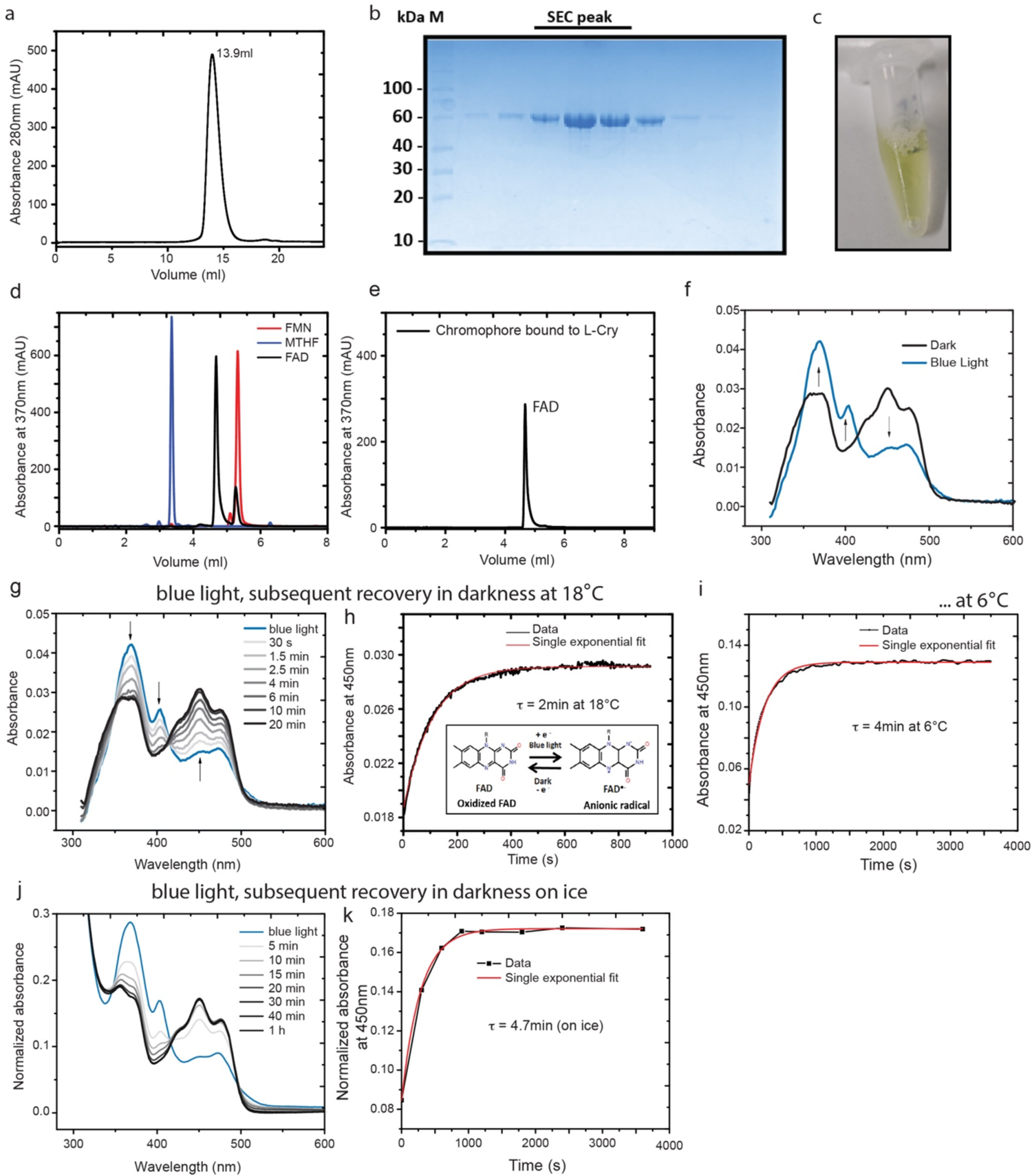
Purification, biochemical and spectral characterization of L-Cry with blue light. **(a)** Size-exclusion chromatography (SEC) of L-Cry on analytical S200 10/300 column. L-Cry elutes at 13.9 ml, suggesting a homodimer based on calibration standards. **(b)** 10% Bis-Tris gel loaded with fractions from the L-Cry SEC peak in (a). **(c)** L-Cry protein solution (5 mg/ml) with yellow color from bound oxidized FAD. **(d,e)** Reverse phase HPLC analysis identifies FAD as only L-Cry chromophore. Elution profile of standard chromophores FMN, MTHF and FAD (d) were compared with the L-Cry bound chromophore obtained after heat denaturation (e). **(f)** Absorption spectrum of L-Cry in dark (black) and after 110s blue light (blue). Arrows indicate the change in absorbance at 370nm, 404nm and 450nm between FAD (dark) and FAD°^-^ (after blue light). **(g,h)** L-Cry dark recovery after blue light activation at 18°C. Full Spectra in (g), 450nm absorbance in (h). Inset in (h): Schematic of FAD photoreaction. **(i)** Dark recovery after blue light activation at 6°C (450 nm absorbance). **(j,k)** Dark recovery after blue light activation on ice. Full spectra in J, 450nm absorbance in k.

**Extended Data Figure 6:**
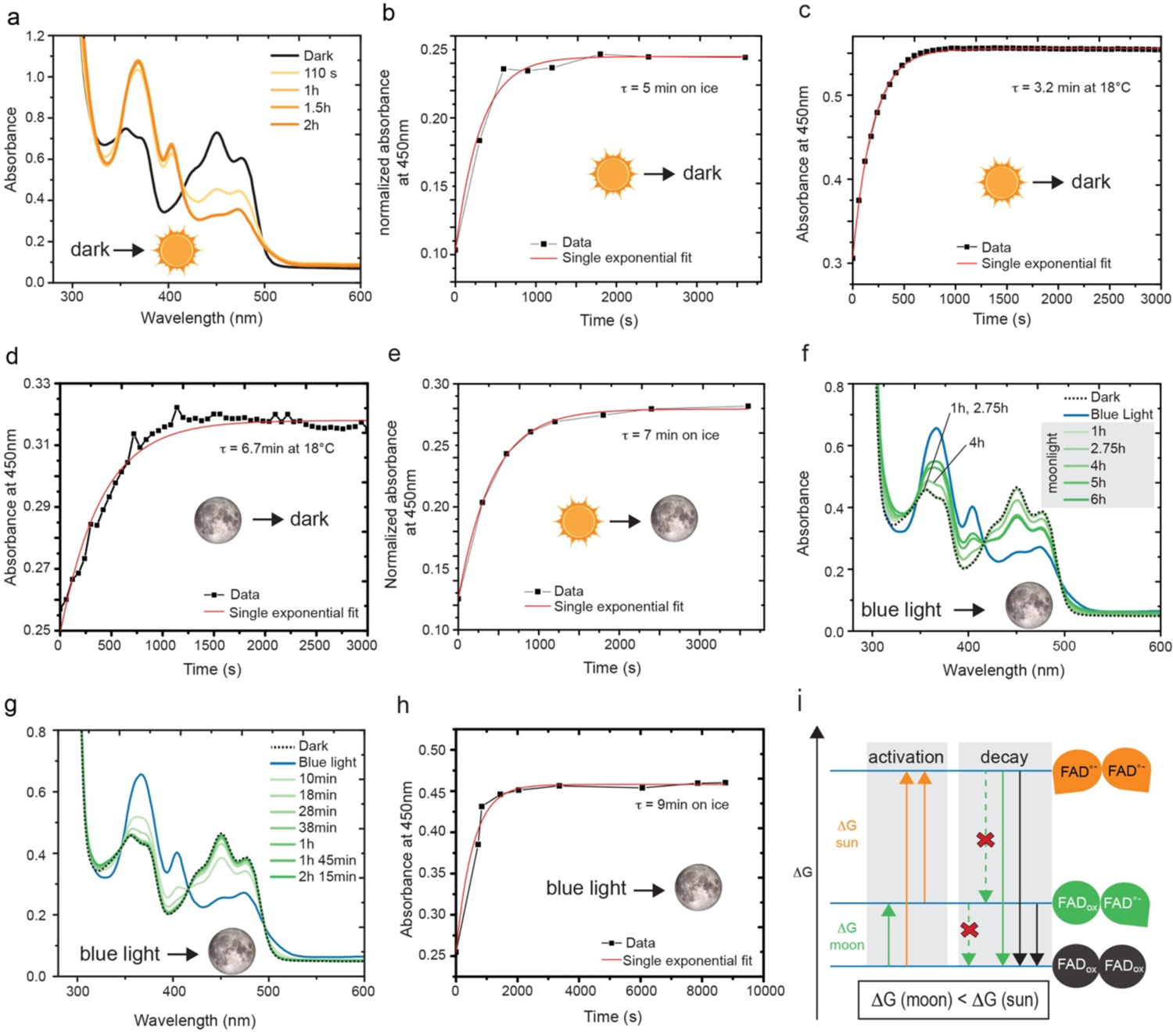
Spectral characterization of L-Cry under naturalistic sun- and moonlight. **(a)** Absorption spectrum of L-Cry in dark (black) and after sunlight exposure (orange). Additional timepoints shown in Fig. 5b. **(b)** Dark recovery of L-Cry after 20min of sunlight on ice: absorbance at 450 nm, full spectra in Fig. 5c. **(c)** Dark recovery of L-Cry after 20min sunlight at 18°C: absorbance at 450 nm. **(d)** Dark recovery of L-Cry after 6h of naturalistic moonlight: absorbance at 450 nm. Full spectra in Fig. 5f. **(e)** Absorbance at 450nm after 20min sunlight followed by dark-state recovery in presence of moonlight. Full spectra in 5h. **(f)** Absorption spectra of L-Cry after 110s blue light illumination followed by an up to 6h exposure to naturalistic moonlight. L-Cry first returns to the dark state (1h, 2.45 h) and after 3h starts to build the moonlight state (4h, 5h, 6h). **(g)** Recovery of oxidized FAD from the blue light-induced anionic FAD°^-^ radical under naturalistic moonlight shows that L-Cry first returns to the dark state. **(h)** Absorbance values at 450 nm from EDF6g. Note: strong blue light results in L-Cry’s sunlight state. **(i)** Schematic model of relative energy levels (ΔG) and transitions of dark-, moonlight- and sunlight states of L-Cry. MALS and SEC analyses of dark-sate L-Cry (Fig.5a, EDF 5a) show that L-Cry forms homodimers. The energy difference between the dark- and moonlight state (ΔG moon) is significantly smaller than the energy required to get from the dark state to the sunlight state or from the moonlight state to the sunlight state (ΔG sun) (ΔG moon and ΔG sun not drawn to scale). This can be explained by an asymmetric L-Cry dimer with different redox potentials of the flavin cofactors and hence different transition energies between FADox and FAD°^-^ in each monomer. *Activation:* Low intensity moonlight (green arrows) can only photoreduce the lower energy flavin within one monomer resulting in the moonlight state, but cannot overcome the larger energy barrier to photoreduce the second flavin molecule. This is, however, possible in presence of sunlight (orange arrows) with much higher intensity, resulting in the fully photoreduced sunlight state. *Decay:* In darkness (black arrows) or in presence of moonlight, the sunlight state directly decays to the dark state, i.e. moonlight does not maintain the sunlight state. Furthermore, the moonlight state does not accumulate upon decay of the sunlight state in presence of moonlight (upper crossed-out dashed green arrow) or in darkness, likely due to the much faster decay kinetics of the sun- and moonlight states (within min) compared to repopulation of the moonlight state (6 hours). In contrast, moonlight is able to maintain existing moonlight-state L-Cry populations, that have accumulated after 6 h moonlight exposure, i.e. moonlight-state L-Cry does not decay to the dark-state in presence of moonlight (lower crossed-out dashed green arrow), but only in complete darkness (black arrow).

**Extended Data Figure 7:**
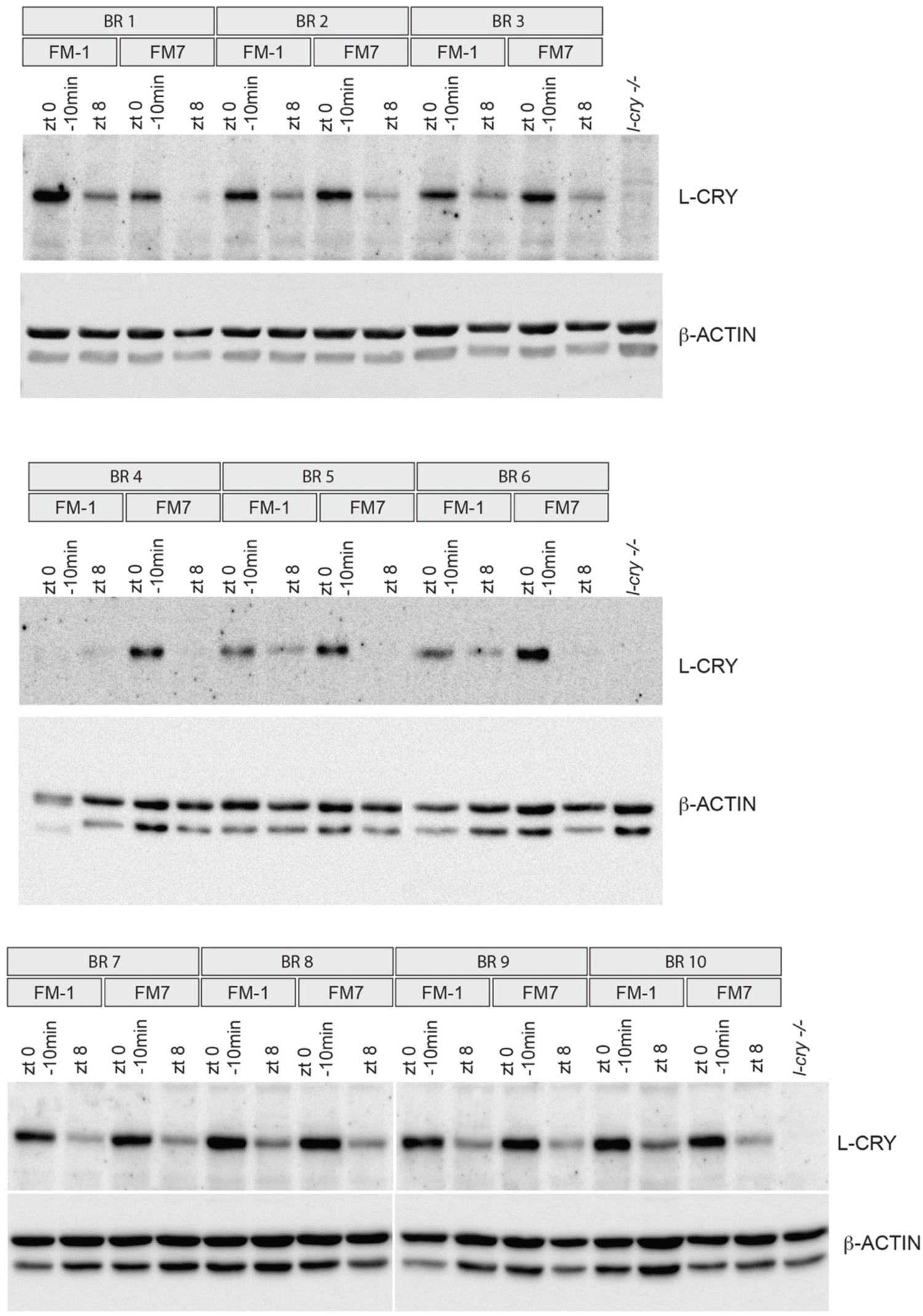
Raw images of all Western blots quantified in **Fig. 6b**. BR: biological replicate.

**Extended Data Figure 8:**
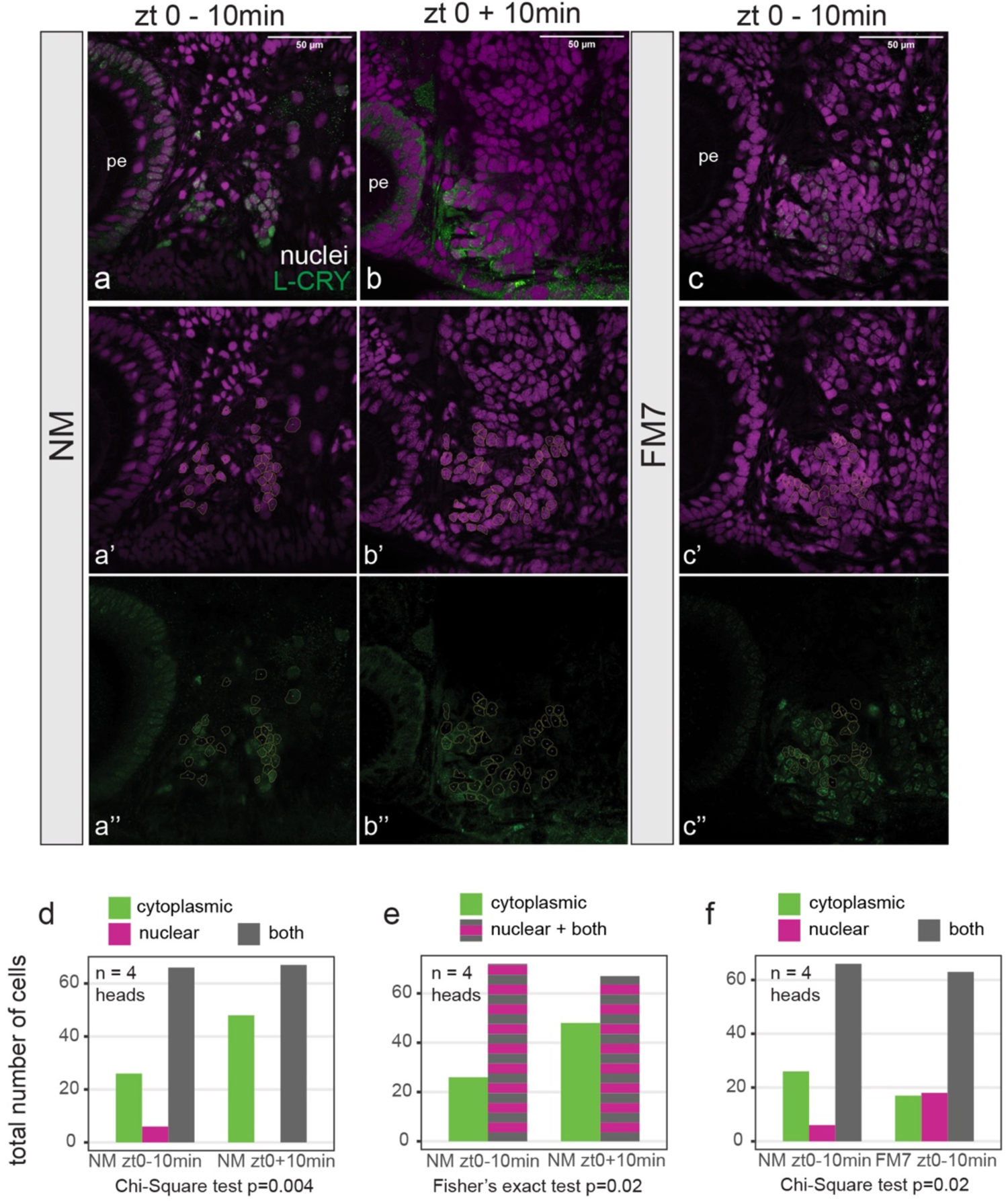
L-Cry immunohistochemistry on *Pdu* heads. **(a-c)** Single layers of confocal images (1.28µm thick) of worm heads stained with anti-L-Cry antibody (green) and HOECHST (magenta) at indicated timepoints. Details of these pictures: Fig. 6e-g’’. pe, posterior eye. **(a’-c’’)** Examples for selection of Regions of Interest (ROI) used for the quantification of L-Cry protein localization in d-f. **(d-f)** Categorical quantification of L-Cry’s subcellular localization at indicated timepoints. The statistical analysis was performed according to the requirements for categorical data (24).

**Extended Data Figure 9:**
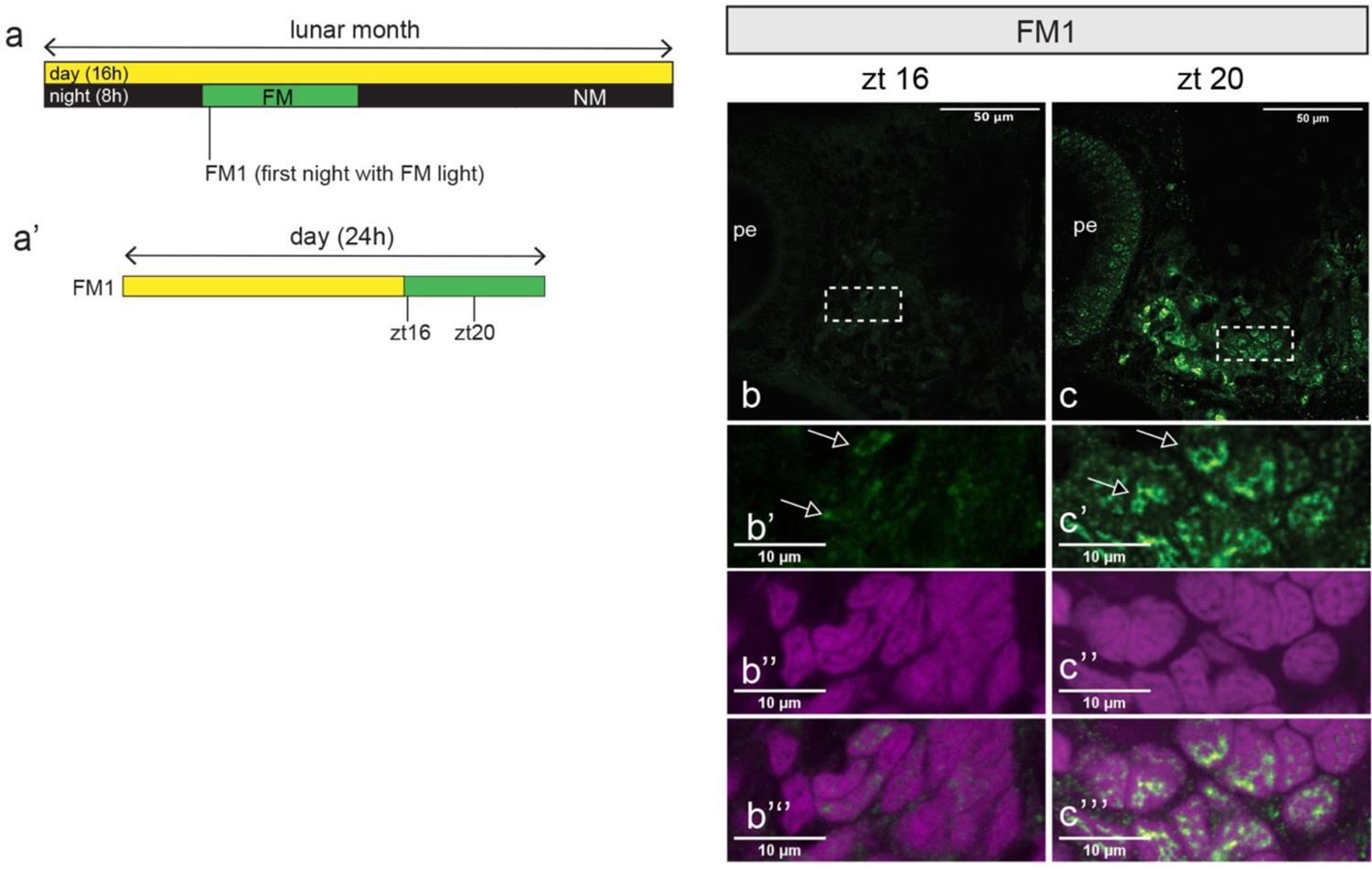
L-Cry protein is endogenously present at timepoints that allow for a sufficiently long exposure to naturalistic moonlight to reach the moonlight-state. **(a,a’)** Scheme of sampling timepoints. 16hrs day (light) and 8 hrs night (dark or moonlight) per 24hrs, with 8 nights of moonlight per month. **(b-c)** Confocal single layer (1.28µm) images of worm heads stained with anti-L-Cry antibody (green). White rectangles: areas of the zoom-ins presented below. **(b*’*-c’’’)** zoomed pictures of the areas depicted in b-c. anti-L-Cry antibody (green), HOECHST (magenta: nuclei), Arrows indicate L-Cry staining. Scale bars: 10um.

## Supplementary Table

**Supplementary Table 1.** Raw data obtained from scoring L-Cry subcellular localization signal (see Extended Data Figure 8) using 40x confocal microscopy images like shown in the example in EDF8. (Excel table)

## References

1. Fox, H. M. Lunar periodicity in Reproduction. Proceedings of the Royal Society of London **Series B** 95, 523–550 (1924).

2. Levy, O. et al. Light-responsive cryptochromes from a simple multicellular animal, the coral *Acropora millepora*. Science 318, 467–470 (2007).

3. Numata, H. & Helm, B. Annual, lunar, and tidal clocks: patterns and mechanisms of nature’s enigmatic rhythms. (Springer, 2014).

4. Korringa, P. Relations between the moon and periodicity in the breeding of marine animals. Ecological Monographs 17, 347–381 (1947).

5. Hauenschild, C. Lunar periodicity. Cold Spring Harb. Symp. Quant. Biol. 25, 491–497 (1960).

6. Tessmar-Raible, K., Raible, F. & Arboleda, E. Another place, another timer: Marine species and the rhythms of life. Bioessays 33, 165–172, doi:10.1002/bies.201000096 (2011).

7. Bézy, V. S. et al. Mass-nesting events in olive ridley sea turtles: environmental predictors of timing and size. Anim. Behav. 163, 85–94, doi:https://doi.org/10.1016/j.anbehav.2020.03.002 (2020).

8. Norevik, G., Akesson, S., Andersson, A., Backman, J. & Hedenstrom, A. The lunar cycle drives migration of a nocturnal bird. PLoS Biol. 17, e3000456, doi:10.1371/journal.pbio.3000456 (2019).

9. Raible, F., Takekata, H. & Tessmar-Raible, K. An Overview of Monthly Rhythms and Clocks. Frontiers in Neurology 8, doi:ARTN 18910.3389/fneur.2017.00189 (2017).

10. Casiraghi, L. et al. Moonstruck sleep: Synchronization of human sleep with the moon cycle under field conditions. Sci Adv 7, doi:10.1126/sciadv.abe0465 (2021).

11. Helfrich-Forster, C. et al. Women temporarily synchronize their menstrual cycles with the luminance and gravimetric cycles of the Moon. Sci Adv 7, doi:10.1126/sciadv.abe1358 (2021).

12. Shlesinger, T. & Loya, Y. Breakdown in spawning synchrony: A silent threat to coral persistence. Science 365, 1002–1007, doi:10.1126/science.aax0110 (2019).

13. Neumann, D. Temperature compensation of circasemilunar timing in the intertidal insect *Clunio*. Journal of Comparative Physiology A - Sensory Neural and Behavioral Physiology 163, 671–676 (1988).

14. Franke, H.-D. The Role of Light and Endogenous Factors in the Timing of the Reproductive Cycle of Typosyllis prolifera and Some Other Polychaetes. Am. Zool. 26, 433–445 (1986).

15. Zantke, J. et al. Circadian and Circalunar Clock Interactions in a Marine Annelid. Cell Reports 5, 99–113, doi:10.1016/j.celrep.2013.08.031 (2013).

16. Franke, H. D. On a clocklike mechanism timing lunar-rhythmic reproduction inTyposyllis prolifera (Polychaeta). *Journal of Comparative Physiology A: Neuroethology, Sensory*, Neural, and Behavioral Physiology 156, 553–561 (1985).

17. Hauenschild, C. Über das lunarperiodische Schwärmen von *Platynereis dumerilii* in Laboratoriumszuchten. Naturwissenschaften 41, 556–557 (1954).

18. Fukunaga, K., Yamashina, F., Takeuchi, Y., Yamauchi, C. & Takemura, A. Moonlight is a key entrainer of lunar clock in the brain of the tropical grouper with full moon preference. BMC Zoology 5, 11, doi:10.1186/s40850-020-00060-8 (2020).

19. Fukushiro, M. et al. Lunar phase-dependent expression of cryptochrome and a photoperiodic mechanism for lunar phase-recognition in a reef fish, goldlined spinefoot. PLoS ONE 6, e28643, doi:10.1371/journal.pone.0028643 (2011).

20. Brady, A. K., Willis, B. L., Harder, L. D. & Vize, P. D. Lunar Phase Modulates Circadian Gene Expression Cycles in the Broadcast Spawning Coral Acropora millepora. Biol. Bull. 230, 130–142, doi:10.1086/BBLv230n2p130 (2016).

21. Andreatta, G. & Tessmar-Raible, K. The Still Dark Side of the Moon: Molecular Mechanisms of Lunar-Controlled Rhythms and Clocks. J. Mol. Biol., doi:10.1016/j.jmb.2020.03.009 (2020).

22. Kaiser, T. S. et al. The genomic basis of circadian and circalunar timing adaptations in a midge. Nature 540, 63–73, doi:10.1038/nature20151 (2016).

23. Bannister, S. et al. TALE Nucleases mediate efficient, heritable genome modifications in the marine annelid *Platynereis dumerilii*. Genetics 197, 19–31, doi:10.1534/genetics.112.148254 (2014).

24. Fisher, N. I. Statistical analysis of circular data. (cambridge university press, 1995).

25. Revilla, I. D. R. et al. Characterization of cephalic and non-cephalic sensory cell types provides insight into joint photo- and mechanoreceptor evolution. Elife 10, doi:10.7554/eLife.66144 (2021).

26. Veedin Rajan, V. B. et al. Seasonally relevant UVA light alters neurohormone amounts and behavior via a ciliary opsin in a marine mass spawning annelid. Nature Ecology & Evolution, doi:10.1038/s41559-020-01356-1 (2021).

27. Ayers, T., Tsukamoto, H., Gühmann, M. & Tessmar-Raible, K. A Go-type Opsin mediates the Shadow Reflex in the annelid Platynereis dumerilii. BMC Biol. 16, 41, doi:10.1186/s12915-018-0505-8 (2018).

28. Guehmann, M. et al. Spectral Tuning of Phototaxis by a Go-Opsin in the Rhabdomeric Eyes of Platynereis. Curr. Biol. 25, 2265–2271, doi:10.1016/j.cub.2015.07.017 (2015).

29. Ranzi, S. Ricerche sulla biologia sessuale degli Anellidi. Pubbl. Staz. Zool. Napoli 11, 271–292 (1931).

30. Zantke, J., Bannister, S., Veedin Rajan, V. B., Raible, F. & Tessmar-Raible, K. Genetic and Genomic Tools for the marine annelid *Platynereis dumerilii*. Genetics 197, 9–31, doi:10.1534/genetics.112.148254 (2014).

31. Ranzi, S. Maturita sessuale degli Anellidi e fasi lunari. Boll. Soc. Ital. Biol. Sperim. 6, 18 (1931).

32. Zurl, M., et al. Two light sensors decode moonlight versus sunlight to adjust a plastic circadian/circalunidian clock to moon phase. bioRxiv https://doi.org/10.1101/2021.04.16.440114 (2021).

33. Berndt, A. et al. A novel photoreaction mechanism for the circadian blue light photoreceptor *Drosophila* cryptochrome. J. Biol. Chem. 282, 13011–13021, doi:M608872200 [pii]10.1074/jbc.M608872200 [doi] (2007).

34. Xu, B., Feng, X. & Burdine, R. D. Categorical data analysis in experimental biology. Dev Biol 348, 3–11, doi:10.1016/j.ydbio.2010.08.018 (2010).

35. Peschel, N., Chen, K. F., Szabo, G. & Stanewsky, R. Light-Dependent Interactions between the Drosophila Circadian Clock Factors Cryptochrome, Jetlag, and Timeless. Curr. Biol. 19, 241–247, doi:10.1016/j.cub.2008.12.042 (2009).

36. Czarna, A. et al. Structures of Drosophila Cryptochrome and Mouse Cryptochrome1 Provide Insight into Circadian Function. Cell 153, 1394–1405, doi:10.1016/j.cell.2013.05.011 (2013).

37. Häfker, N. S. & Tessmar-Raible, K. Rhythms of behavior: are the times changin’? Curr. Opin. Neurobiol. 60, 55–66, doi:10.1016/j.conb.2019.10.005 (2020).

38. Oliveri, P. et al. The Cryptochrome/Photolyase Family in aquatic organisms. Marine Genomics 14, 23–37, doi:10.1016/j.margen.2014.02.001 (2014).

39. Hauenschild, C. & Fischer, A. Platynereis dumerilii. Mikroskopische Anatomie, Fortpflanzung, Entwicklung. [Platynereis dumerilii. Microscopical anatomy, reproduction and development] 1-55 (Stuttgart, 1969).

40. Tessmar-Raible, K., Steinmetz, P. R., Snyman, H., Hassel, M. & Arendt, D. Fluorescent two-color whole mount in situ hybridization in Platynereis dumerilii (Polychaeta, Annelida), an emerging marine molecular model for evolution and development. Biotechniques 39, 460, 462, 464 (2005).

41. Schindelin, J. et al. Fiji: an open-source platform for biological-image analysis. Nat. Methods 9, 676–682, doi:10.1038/nmeth.2019 (2012).

42. Stringer, C., Wang, T., Michaelos, M. & Pachitariu, M. Cellpose: a generalist algorithm for cellular segmentation. Nat. Methods 18, 100–106, doi:10.1038/s41592-020-01018-x (2021).

43. Huang, R. S., Chen, C. F. & Sereno, M. I. Mapping the complex topological organization of the human parietal face area. Neuroimage 163, 459–470, doi:10.1016/j.neuroimage.2017.09.004 (2017).

44. Lee, A. Circular data. WIREs Computational Statistics 2, 477–486, doi: https://doi.org/10.1002/wics.98 (2010).

45. Landler, L., Ruxton, G. D. & Malkemper, E. P. Circular data in biology: advice for effectively implementing statistical procedures. Behav. Ecol. Sociobiol. 72, 128, doi:10.1007/s00265-018-2538-y (2018).

46. Backfisch, B. et al. Stable transgenesis in the marine annelid *Platynereis dumerilii* sheds new light on photoreceptor evolution. Proc Natl Acad Sci USA 110, 193–198, doi:10.1073/pnas.1209657109 (2013).

47. R: A Language and Environment for Statistical Computing. (https://www.R-project.org/ Vienna, Austria, 2016).

